# m^6^A RNA methylation modulates Zika virus infection by regulating serine proteases in *Aedes albopictus*

**DOI:** 10.64898/2026.03.16.711780

**Authors:** Anderson de Mendonça Amarante, Lucas Tirloni, Jonas Koch, Felix Bormann, Serena Rosignoli, Alessandro Paiardini, Dante Rotili, Tarcisio Brito, Laura de Andrade Tavares, Juan Diego de Paula Li Yasumura, Vitor Coutinho Carneiro, Laura Menendez Kury, Attilio Pane, Frank Lyko, Marcelo Rosado Fantappié

## Abstract

Epitranscriptomic RNA modifications, particularly N^6^-methyladenosine (m⁶A), have emerged as important regulators of host–virus interctions. However, the role of m⁶A in arbovirus infection within mosquito vectors remains poorly defined. Here, we characterized the m⁶A RNA methylation machinery in *Aedes albopictus* C6/36 cells and examined its contribution to Zika virus (ZIKV) replication. Arbovirus infection did not significantly alter the transcriptional levels or enzymatic activity of the core m⁶A methyltransferase components METTL3 and METTL14. In contrast, pharmacological inhibition of METTL3 markedly enhanced ZIKV replication, indicating an antiviral role for m⁶A in mosquito cells. Transcriptome-wide analysis of C6/36 cells treated with the METTL3 inhibitor STM2457 revealed extensive changes in gene expression, including the pronounced upregulation of multiple serine proteases, particularly members of the CLIP family. Single-nucleotide–resolution mapping of m⁶A using GLORI-sequencing showed that m^6^A is absent from Zika virus RNA, but readily detectable in the *A. albopictus* transcriptome. Data analysis defined key features of the mosquito epitranscriptome and demonstrated that m⁶A modifications are enriched within the coding regions of serine protease transcripts, supporting their direct regulation by m⁶A. Functionally, inhibition of serine protease activity using AEBSF resulted in a significant reduction of ZIKV replication. Together, these findings identify m⁶A RNA methylation as a critical regulator of ZIKV infection in mosquito cells and uncover an epitranscriptomic pathway linking m⁶A-dependent control of serine proteases to vector–virus interactions.

**Author Summary:** Mosquito-borne viruses such as Zika virus pose a major threat to global public health. Successful transmission of these viruses depends not only on infection in humans, but also on their ability to replicate efficiently inside mosquito vectors. Chemical modifications of RNA, collectively known as epitranscriptomic marks, have recently emerged as important regulators of gene expression and virus–host interactions. Among these, N^6^-methyladenosine (m^6^A) is the most abundant internal RNA modification in eukaryotic cells. While m^6^A has been extensively studied in mammalian systems, its role in mosquito antiviral responses remains poorly understood. In this study, we investigated how m^6^A RNA methylation influences Zika virus infection in mosquito cells derived from *Aedes albopictus*. We found that reducing m^6^A levels enhances viral replication, indicating that this RNA modification restricts infection in mosquito cells. Notably, Zika virus RNA itself does not contain detectable m^6^A modifications. Instead, m^6^A regulates the expression of specific mosquito genes, including a group of serine proteases that influence viral replication. Pharmacological inhibition of these proteases significantly impaired virus growth, identifying them as key downstream effectors. Our findings reveal an antiviral role for m^6^A in mosquito cells and uncover a previously unrecognized epitranscriptomic pathway that shapes mosquito–virus interactions. Understanding how RNA modifications regulate arbovirus infection in vectors may open new avenues for strategies aimed at limiting virus transmission.

## Introduction

Viruses are obligate intracellular agents that depend on host cells to replicate and spread. To create a favorable environment for their replication, viruses often hijack and remodel host molecular pathways, including those involved in gene regulation. A growing body of evidence indicates that viral infections can profoundly impact host gene expression programs, leading to modifications in chromatin accessibility, histone post-translational modifications, DNA methylation, and RNA modifications [1–3]. These epigenetic changes not only support viral genome expression and replication but also modulate cellular antiviral defenses, thereby shaping the outcome of infection [3,4]. Epigenetic and epitranscriptomic reprogramming are now recognized as a key mechanism by which viruses establish infection, persist within the host, or evade immune surveillance [5–8].

Among the diverse epitranscriptomic regulators, N^6^-methyladenosine (m^6^A) RNA methylation represents the most prevalent internal modification of eukaryotic messenger RNAs and noncoding RNAs [9–11]. m^6^A is installed by a multiprotein methyltransferase complex known as the “writers” (including METTL3, METTL14, and accessory factors), removed by demethylases termed “erasers” (such as FTO and ALKBH5), and recognized by effector proteins referred to as “readers” (notably the YTH domain-containing family) [10,12,13]. Through this dynamic and reversible machinery, m^6^A exerts broad control over RNA metabolism, including alternative splicing, nuclear export, transcript stability, and translational efficiency [12] Consequently, m^6^A serves as a versatile regulatory layer that fine-tunes gene expression in processes ranging from embryogenesis and tissue differentiation to stress responses and immune signaling [12,14–16]. However, despite a growing body of work on viruses such as HIV [17], influenza [18], and hepatitis C [19], relatively few studies have specifically addressed the role of m^6^A in arboviruses [3,20–22], a diverse group of medically important viruses transmitted by arthropod vectors.

The available literature suggests that m^6^A can exert both proviral and antiviral effects in arbovirus infections, depending on the virus species and the cellular context [3,20–22]. For example, studies in flaviviruses, including Dengue and Zika, have indicated that m^6^A modifications may regulate viral RNA stability and replication efficiency [20–22]. Conversely, in other systems, depletion of m^6^A writers has been associated with enhanced viral replication, highlighting the complexity and context-dependence of this modification [23].

A central point of debate in the field concerns whether arbovirus genomes themselves are directly modified by m^6^A. While some studies employing m^6^A-specific antibodies and high-throughput sequencing have mapped putative methylation sites within viral RNA [3,20–22], a recent publication has questioned the specificity and reproducibility of these findings, suggesting that additional technical rigor and orthogonal validation are required [24]. Thus, the extent to which arbovirus genomes are bona fide substrates for m^6^A methylation remains unresolved, and the functional implications of such modifications are still under active investigation.

Elucidating the interplay between host epitranscriptomic regulation and viral replication is essential for a more comprehensive understanding of arbovirus biology. In this context, the mosquito host represents a particularly relevant system, as it serves not only as the vector that transmits the virus to humans but also as a biological environment where viruses must overcome multiple cellular barriers in order to replicate and persist. Unlike mammalian hosts, mosquitoes maintain long-term, largely asymptomatic viral infections, suggesting that finely tuned molecular mechanisms regulate the balance between viral replication and host survival. Identifying the mosquito molecular players that mediate these processes—especially those related to epitranscriptomic regulation may therefore uncover novel aspects of virus-vector interactions. Such insights could provide new opportunities for the development of antiviral strategies that exploit mosquito-specific pathways, including those governed by RNA modifications such as m^6^A methylation, with the potential to disrupt transmission at its source.

## Materials and methods

### Sequence analysis, protein alignment, and primer design

Nucleotide and amino acid sequences corresponding to components of the m^6^A RNA methylation machinery (IDs listed in the supplementary Table 1) from *Aedes aegypti* (*A.ae*) and *Aedes albopictus* (*A.al*) were retrieved using the online BLASTp and BLASTn tools (blast.ncbi.nlm.nih.gov/Blast). Searches were performed using orthologous sequences from *Drosophila melanogaster* and *Homo sapiens* as queries.

Identified sequences were selected based on the following criteria: E-value < 1 × 10⁻⁵, sequence coverage > 45%, and amino acid identity > 30% [25]. Multiple sequence alignments of amino acid sequences were performed using ClustalW, and functional domains were identified using the ExPASy-PROSITE tool (prosite.expasy.org) [26].

Primers used for RT-qPCR analyses and double-stranded RNA (dsRNA) synthesis were designed using Primer-BLAST (www.ncbi.nlm.nih.gov/tools/primer-blast).

### Cell lines and viruses

The Vero (“African green monkey kidney”) cell line, C6/36 (*A. albopictus* embryonic cells) [27], and Aag2 (*A. aegypti* embryonic cells) [28] were used in this study. Vero cells were maintained in Dulbecco’s Modified Eagle Medium (DMEM) (Invitrogen, USA) at 37 °C in a humidified atmosphere containing 5% CO₂. In contrast, C6/36 cells were cultured in Leibovitz’s L-15 medium (Invitrogen, USA), and Aag2 cells were cultured in Schneider’s Drosophila medium (Invitrogen, USA); both mosquito cell lines were maintained at 28 °C without CO₂ supplementation.

All culture media were supplemented with 10% fetal bovine serum (Gibco, USA), 100 U/mL penicillin, and 100 µg/mL streptomycin (Merck). Zika virus (ZIKV; strain ZIKV/H.sapiens/Brazil/PE243/2015) and Chikungunya virus (CHIKV; strain S27-African) were propagated in C6/36 cells for 6 days and viral stocks were stored at −80 °C.

### Inhibitor treatments and viral infection

C6/36 cells were seeded in 12-well culture plates and grown to approximately 50% confluence prior to treatment. Cells were pretreated with the METTL3 inhibitor STM2457 [29] at concentrations of 20 or 40 µM for 48 h, while control cells were treated with an equivalent volume of DMSO (Merck). Treatment efficiency was assessed by immunodetection using dot blot analysis and by mass spectrometry. Following pretreatment, cells were infected with ZIKV at a multiplicity of infection (MOI) of 1.0 and maintained for 3 days post-infection. Viral loads were quantified by plaque assay and by RT-qPCR.

For experiments involving the serine protease inhibitor AEBSF (Merck), C6/36 cells were seeded in 12-well plates and pretreated at approximately 50% confluence with AEBSF at concentrations of 200, 300, or 400 µM for 24 h. Cells pretreated with 300 µM AEBSF were subsequently subjected to viral binding and internalization assays. Following pretreatment with either STM2457 or AEBSF, cell viability was determined using the CellTiter-Glo assay according to the manufacturer’s instructions (Promega, USA).

### Plaque assay

Vero cells were seeded in 24-well plates and grown to 80–90% confluence. Zika virus samples were serially diluted (from 10⁻¹ to 10⁻⁶), and 200 µL of each dilution was inoculated into individual wells. Viral inocula were allowed to adsorb for 1 h at 37 °C, after which the inoculum was removed and cells were overlaid with 500 µL of DMEM containing 1% carboxymethylcellulose (Sigma), supplemented with 2% fetal bovine serum (Gibco, USA), 100 U/mL penicillin, and 100 µg/mL streptomycin (Merck).

At 3 to 5 days post-infection, cells were fixed with 4% formaldehyde for 1 h, washed with distilled water, and stained with 1% crystal violet solution for 15 min. Plaques were counted, and viral titers were calculated and expressed as plaque-forming units per milliliter (PFU/mL), as previously described [30].

### RNA extraction and RT-qPCR

Total cellular RNA was isolated using TRIzol reagent (Thermo Fisher Scientific, USA) according to the manufacturer’s instructions. Subsequently, 1 µg of total RNA was treated with DNase (Ambion) and reverse-transcribed using the High-Capacity cDNA Reverse Transcription Kit (Thermo Fisher Scientific, USA) following the manufacturer’s protocol.

For gene expression analyses, quantitative PCR was performed using the StepOnePlus Real-Time PCR System (Applied Biosystems) and GoTaq® qPCR Master Mix (Promega). Relative mRNA abundance was determined using the comparative Ct (ΔΔCt) method [31], with the ribosomal protein gene RP49 used as an endogenous control [32]. All primers used in this study is listed in the supplementary Table 2.

### Dot blot analysis

For immunodetection of m^6^A RNA modification, 500 ng of total RNA were spotted onto PVDF membranes and UV-crosslinked at 80 °C for 40 min. Membranes were blocked for 1 h in blocking buffer (1× PBS, 0.05% Tween-20, and 3% bovine serum albumin) and incubated overnight (16 h) at 4 °C with a monoclonal anti-m6A primary antibody (Thermo Fisher Scientific). Membranes were then washed with 1× PBS containing 0.05% Tween-20 and incubated for 1 h with an HRP-conjugated anti-rabbit secondary antibody (Thermo Fisher Scientific). Immunoreactive signals were detected by chemiluminescence using SuperSignal West Dura substrate (Thermo Fisher Scientific) and imaged with an Amersham Imager 600 system (GE Healthcare).

For RNA loading control, 500 ng of total RNA were spotted onto PVDF membranes, UV-crosslinked at 80 °C for 40 min, and stained with 0.04% methylene blue solution (Sigma). Signal intensities were quantified by densitometric analysis using ImageJ software, and m6A signals were normalized to their corresponding methylene blue loading controls.

### Quantification of m^6^A Levels in Total RNA by LC–MS/MS

LC-MS/MS analysis of total RNA was performed as described before [33].

### RNA interference and viral infection

For RNA interference (RNAi) experiments, double-stranded RNAs (dsRNAs) were synthesized using the T7 MEGAscript in vitro transcription kit (Thermo Fisher Scientific, USA) according to the manufacturer’s instructions. Templates corresponding to target genes were generated by PCR using gene-specific primers (listed in the supplementary table), in which 2 µL of cDNA derived from Aag2 cells were amplified using PrimeSTAR Max DNA Polymerase (Takara). Control dsRNA (dsLuciferase) was generated from the luciferase gene template, amplified under the same conditions using 10 ng of the pGL4.10 Luciferase Reporter plasmid.

For transfection, 2 × 10^5^ Aag2 cells were seeded in 12-well plates and cultured for 24 h. Subsequently, 1 µg of dsRNA per well was transfected using Attractene reagent (Qiagen) following the manufacturer’s instructions. After 48 h, cells were transfected a second time under the same conditions. Forty-eight hours after the second transfection, cells were trypsinized and replated into 12-well plates at a density of 2 × 10^5^ cells per well and cultured for an additional 24 h. Cells were then infected with ZIKV at a multiplicity of infection (MOI) of 1.0 for 3 days. Viral loads were subsequently quantified by plaque assay and by RT-qPCR.

### Immunofluorescence microscopy

Approximately 4 × 10^5^ cells were seeded onto glass coverslips placed in 24-well plates. Cells were fixed with 4% paraformaldehyde for 1 h and permeabilized for 15 min with 1× PBS containing 0.15% Triton X-100. Cells were then blocked for 1 h with blocking buffer (1× PBS, 0.05% Tween-20, and 10% bovine serum albumin). Following blocking, cells were incubated overnight (16 h) with an anti-m6A primary antibody.

Cells were subsequently washed with 1× PBS containing 0.1% Tween-20 and incubated for 1 h with an Alexa Fluor 488–conjugated anti-rabbit secondary antibody. After additional washes, coverslips were mounted using ProLong Gold Antifade Mountant with DAPI (Thermo Fisher Scientific). Immunofluorescence images were acquired using a Zeiss LSM 710 confocal microscope.

### Transcriptomic analysis

#### Library preparation, sequencing, and data analysis

Total RNA was isolated using the TRIzol^TM^ Reagent (Invitrogen) according to the manufacturer’s instructions. RNA integrity and quantity were assessed using a Bioanalyzer system (Agilent Technologies). The Illumina libraries were constructed using the NEBNextUltraTM II (Directional) RNA with polyA selection library prep kit and sequencing was performed in an Illumina Novaseq 6000 DNA sequencer (Novogene). RNA-Seq reads underwent quality control, adapter trimming, mapping, counting and differential expression analysis via the mRNAseq pipeline from the snakePipes package (v2.7.3) [34]. Mapping was performed using the software STAR (v2.7.10b) [35] and the AalbF5 (GCF_035046485) version of the *A. albopictus* genome as reference. Read count for each gene was done using featureCounts (v2.0.1) [36] and differential expression with the R package DESeq2 (v1.38.3) [37]. Variance-stabilizing transformation was used for downstream analyses and data visualization. Thresholds for differentially expressed genes were set to adjusted p-value < 0.01 and absolute log2 fold change > 0.5. Functional analyses were performed using topGO package [38] with the GO Association File (GAF) from the AalbF5 annotation, applying the *elim* algorithm, Fisher’s exact test and only considering terms with p-value < 0.05.

### Molecular Docking

#### Structural Modeling of METTL3/METTL14 and Non-covalent Docking

To model the *A. aegypti* METTL3/METTL14 heterodimer, the high-resolution human METTL3–METTL14 crystal structure (PDB ID: 7O2I) [39] [36] was used as a structural reference. Protein sequences were aligned to their human orthologs using MUSCLE [40], confirming strong conservation across the methyltransferase (MTase) domains. Structural prediction of the *A. albopictus* heterodimer was performed using AlphaFold2 [41], guided by the human heterodimer to preserve domain orientation and catalytic site geometry. Particular attention was given to alignment of the SAM-binding pocket and active-site residues.

Protein structures were prepared using AutoDockTools [42], with addition of polar hydrogens, assignment of Gasteiger charges, and optimization for docking. The METTL3 inhibitor STM2457 was modeled based on published structures; ligand coordinates were generated, protonated at physiological pH, and charged accordingly. Docking simulations were performed using AutoDock Vina [43], with the search grid centered on the METTL3 SAM-binding pocket and encompassing adjacent binding regions. Default exhaustiveness parameters were used, and top-scoring poses were visually inspected to confirm productive active-site engagement.

For validation, the docked STM2457 pose was superimposed onto the human METTL3/METTL14–STM2457 crystal structure using PyMOL [44], allowing direct comparison of binding orientation and conserved residue interactions.

#### Protein Structure Prediction for AEBSF Covalent Docking Targets

All protease and viral protein sequences used for AEBSF docking were retrieved from UniProt [45]. Three-dimensional structures were predicted using AlphaFold3 with standard parameters (five recycles, no structural templates, AMBER relaxation enabled). Structural models were selected based on high per-residue confidence scores (pLDDT > 80). Predicted alignment error (PAE) maps were examined to confirm intra-domain stability, with acceptable models showing intra-domain errors below 10 Å. For enzymes with known human homologs, active-site residues were identified based on conserved catalytic motifs and structural homology to experimentally solved structures deposited in the RCSB Protein Data Bank.

#### Covalent docking, model refinement and interaction analysis

Protein structures were prepared for docking using AutoDockTools (ADT v1.5.7) [43] by adding polar hydrogens, assigning Kollman charges, and removing crystallographic water molecules and non-essential ions. Structures were converted to PDBQT format with defined partial charges and rotatable bonds.

Covalent docking of AEBSF was performed using AutoDockFR (ADFR), implemented through DockingPie in PyMOL. Docking parameters included an exhaustiveness of 32, the Lamarckian genetic algorithm, 256 independent runs, and 25 million energy evaluations. The top ten ligand poses were clustered based on RMSD (< 2.0 Å) and ranked according to predicted binding free energy (ΔG, kcal/mol). Docked complexes were refined via energy minimization using GROMACS (version 2023.1) employing the steepest descent algorithm (5,000 steps). Structures were protonated at pH 7.0, and atomic charges were assigned using the AMBER19 force field. Final complexes were analyzed in PyMOL to assess hydrogen bonding, hydrophobic contacts, steric complementarity, and covalent linkage geometry.

#### GLORI-seq library preparation

GLORI-seq library preparation was performed as described previously [46]. Briefly, total RNA was purified by ethanol precipitation and then fragmented by incubation for 3 min at 94 °C using the NEBNext Magnesium RNA Fragmentation Module (New England Biolabs). Fragmented total RNA was then DNase I-treated and purified by the RNA Clean & Concentrator-5 kit (Zymo Research). RNA protection, deamination, and deprotection were performed as described in Liu et al., 2023; Shen et al., 2024, using 200 ng of fragmented total RNA as input material. RNA samples were supplemented with a synthetic Spike-In RNA oligonucleotide (5’-AGCGCAGCGAACAUGACACGUGCUCAACAGUGUAGUUGGACUUCCUCGCAUCUAGCCGCUAUGCACGUGCAGCU-3’).

For the library preparation, a two-step RNA end-repair protocol was applied. RNA 3′-end dephosphorylation was carried out using Antarctic Phosphatase (New England Biolabs). Subsequently, RNA 5′-end phosphorylation was performed using T4 Polynucleotide Kinase (New England Biolabs), and the RNA was purified with the RNA Clean & Concentrator-5 kit (Zymo Research). Next, GLORI-seq libraries were prepared using the NEBNext Small RNA Library Prep Set for Illumina with the NEBNext Multiplex Oligos for Illumina (New England Biolabs). Libraries were size-selected via 8% TBE-PAGE (Thermo Fisher Scientific), isolating fragments between 160 bp and 250 bp. Sequencing was performed on a NovaSeq 6000 platform (Illumina) with a 100 bp paired-end protocol at the Next Generation Sequencing Core Facility of the German Cancer Research Center (DKFZ), Heidelberg.

#### GLORI-seq data analysis

GLORI-seq and analysis of GLORI-seq data were performed as described previously [46]. Briefly, sequencing reads were trimmed using Trim Galore (version 0.6.6) and further processed by the GLORI-tools pipeline as described [[39]. GLORI-tools is available on GitHub: https://github.com/liucongcas/GLORI-tools. m^6^A site selection parameters were set as follows: ≥ 5 variant nucleotides, ≥ 15 coverage of A and G bases, ≥ 0.1 A rate for cells or ≥ 0.01 for viruses. GLORI-tools was executed using: Python 3.8.3, samtools 1.10, STAR 2.7.5c and bowtie 1.3.1. The C6/36 genome and transcriptome reference files (canu_80X_arrow2.2) were downloaded from NCBI (https://ftp.ncbi.nlm.nih.gov/genomes/all/GCF/001/876/365/GCF_001876365.2_canu _80X_arrow2.2/). The genome fasta file was then modified to ensure compatibility with the GLORI-pipeline. In addition, the spike-in sequence and the viral genome (ZIKV or CHIKV) was appended to get the final reference fasta files for read alignment and downstream analyses. GLORI was executed using the standard pipeline settings for identification of m^6^A sites in annotated genes. The GLORI-pipeline output was then used to calculate conversion rates and identify the m^6^A-methylation consensus sequences. In brief, median conversion rates were calculated from all positions in the referbase.mpi output file, corresponding to reference As on the plus strand or Ts on the minus strand respectively with a minimum coverage of 15. M6A-methylation consensus sequences analysis was performed using HOMER 4.11 Significant m^6^A sites were extended by +- 2 nucleotides and analyzed using the parameters -len 5 and -noknown

#### Viral binding and internalization assays

To assess viral binding, C6/36 cells pretreated with 300 µM AEBSF for 24 h were incubated at 4 °C for 30 min prior to infection. Cells were then inoculated with ZIKV at a multiplicity of infection (MOI) of 1.0 and maintained at 4 °C for 1 h to allow viral attachment. Following incubation, cells were washed three times with 1× PBS to remove unbound viral particles. The amount of cell-associated viral RNA was then quantified by RT-qPCR as a measure of viral binding.

For viral internalization assays, following the binding step, viral inocula were removed and replaced with fresh culture medium, and cells were subsequently incubated at 37 °C for an additional 2 h to allow viral entry. Cells were then washed with citrate buffer (50 mM citric acid, 50 mM sodium citrate, pH 3.0) to remove extracellular viral particles. Internalized viral RNA levels were quantified by RT-qPCR.

### Statistical analyses

All statistical analyses were performed using GraphPad Prism software (Prism version 6.0; GraphPad Software, Inc., La Jolla, CA). Student’s *t*-test was applied to determine statistical significance between two groups. Asterisks indicate levels of statistical significance as follows: *p* < 0.05 (\**), p < 0.01 (**), p < 0.001 (\*\*\**); ns, not significant.

## Results

### The m^6^A RNA methylation machinery is conserved in *Aedes* mosquito genomes

Given the still limited number of epigenetic studies in mosquitoes, we sought to systematically investigate whether the canonical m^6^A RNA methylation machinery is encoded in the genomes of *A. aegypti* and *A. albopictus* (Figure 1 and Supplementary Figure 1). Through comprehensive *in silico* analyses, we identified the core components of the m^6^A RNA methylation machinery in both *A. albopictus* and *A. aegypti*, including the writer complex proteins METTL3, METTL14, and WTAP, as well as the major m^6^A reader proteins YTHDC and YTHDF (Figure 1A and B). In contrast, no homologues of the canonical m^6^A eraser proteins FTO or ALKBH5 were detected in either *Aedes* genome despite extensive homology searches. Notably, m^6^A erasers have also not been identified in *Drosophila melanogaster [15]* suggesting that the apparent absence of canonical demethylases may be a conserved feature among dipteran insects.

**Figure 1.**
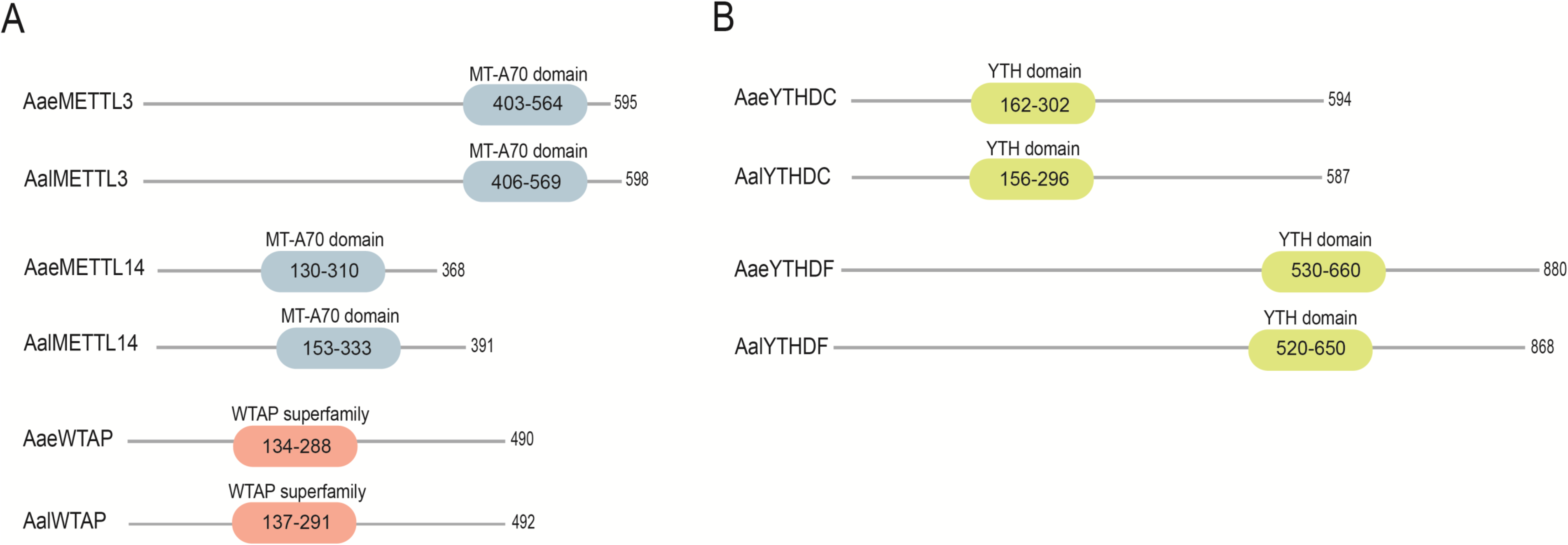
Conserved domain architecture of the m^6^A RNA methylation machinery in *Aedes aegypti* (*A.ae*) and *Aedes albopictus* (*A.al*). Schematic representations of the major components of the m^6^A pathway are shown for both mosquito species, with functional domains indicated by color-coded boxes. Domain annotations were based on conserved motif and sequence analyses. Gene accession numbers corresponding to each protein are listed in Supplementary Table 2. A comparative analysis of full-length amino acid sequences, including functional domain conservations (in black and gray) among *A. aegypti*, *A. albopictus*, *Drosophila melanogaster*, and human orthologs, is provided in Supplementary Figure 1A–E.

Comparative sequence analyses revealed that all identified proteins are encoded as full-length orthologs in both *Aedes* species. Multiple sequence alignments demonstrated a high degree of amino acid identity and similarity between *A. aegypti* and *A. albopictus* across the entire protein length (Supplementary Figure 1A-E). More specifically, a 95% amino acid identity between *A. aegypti* and *A. albopictus* METTL3 proteins was observed (Supplementary Figure 1A-E). These components also exhibited clear sequence homology to their counterparts in other metazoans (Supplementary Figure 1A-E).

Together, these findings indicate that *Aedes* mosquitoes harbor a fully conserved and potentially functional m^6^A RNA methylation machinery, supporting the hypothesis that m^6^A-dependent post-transcriptional regulation is an evolutionarily conserved mechanism in mosquitoes.

### Lack of m^6^A RNA modulation by arbovirus infections

Virus infections are well known to modulate host cellular pathways by altering the expression and/or activity of key regulatory components [1–9]. To determine whether arbovirus infection affects the m^6^A RNA methylation machinery in mosquito cells, we used RT-qPCR to examine the impact of ZIKV infections on the mRNA expression and activity of m^6^A-related enzymes in the *A. albopictus* C6/36 cell line.

Our analyses revealed that infection with ZIKV did not induce detectable changes in the mRNA levels of the core m^6^A writer and reader components (Figure 2A), nor did it alter their functional activity (Figure 2B–C), indicating that arbovirus replication in C6/36 cells does not overtly modulate global activity of the host m^6^A RNA methylation machinery under the conditions tested.

**Figure 2.**
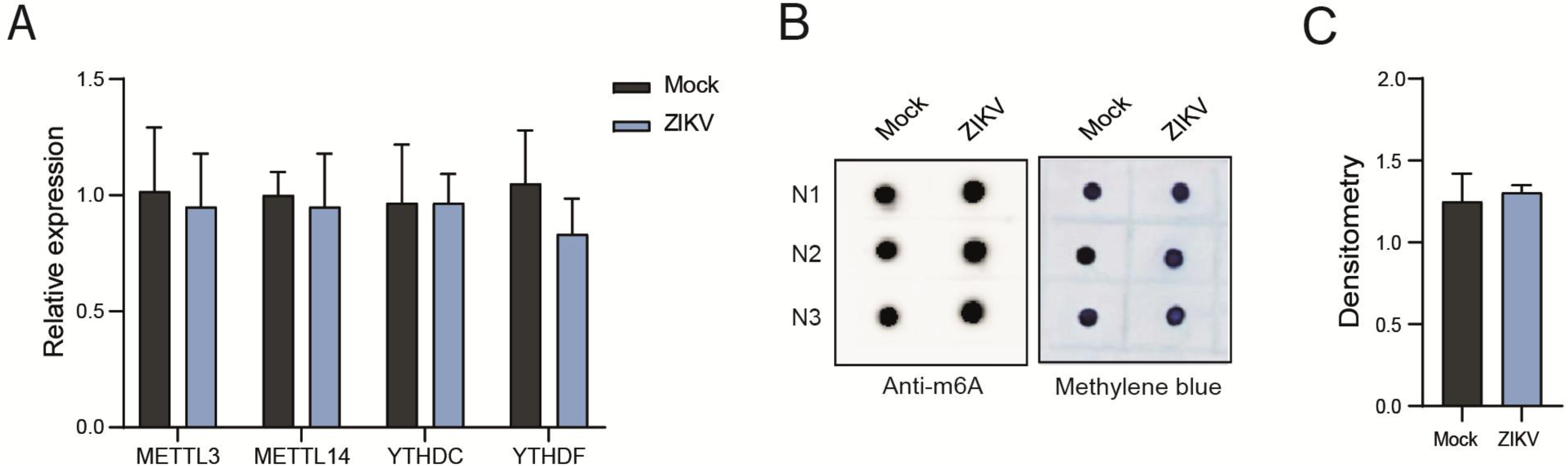
Lack of modulation of the m^6^A pathway following ZIKV infection in C6/36 cells. (A) *A. albopictus* C6/36 cells were either mock-treated or infected with ZIKV at a multiplicity of infection (MOI) of 1.0 and maintained for 3 days post-infection. Total RNA was extracted and the relative mRNA expression levels of core m^6^A machinery components, including METTL3, METTL14, YTHDC, and YTHDF, were quantified by qRT-PCR using gene-specific primers. (B) Total RNA isolated from mock-, or ZIKV-infected C6/36 cells was subjected to dot blot analysis using a monoclonal antibody specific for m6A to assess global m6A RNA levels. Methylene blue staining of the membrane was performed as a loading control to verify equal RNA input. (C) Quantitative densitometric analysis of m^6^A dot blot signal intensities shown in panel B. m^6^A signals were normalized to methylene blue staining and expressed relative to mock-infected controls.

### Molecular docking of STM2457 into the *A. aegypti* METTL3/METTL14 complex

STM2457 is a widely used small-molecule inhibitor of METTL3 complex [29]. To predict the potential inhibitory mechanism of STM2457 on the mosquito m^6^A methyltransferase activity, we performed molecular docking using an AlphaFold-predicted structural model of the *A. aegypti* METTL3/METTL14 complex. The analysis revealed that STM2457 fits into the conserved S-adenosylmethionine (SAM)–binding pocket of the AaMETTL3 subunit, which constitutes the catalytic core of the heterodimer (Figure 3A and B). This predicted binding pose suggests a competitive mode of inhibition, in which STM2457 directly occupies the SAM site and thereby prevents methyl group transfer required for m6A deposition (Figure 3B). The binding orientation of STM2457 in *A. aegypti* METTL3 closely resembles that observed in the human METTL3/METTL14 crystal structure (PDB 7O2I) (Figure 3C) and the docked ligand engages in a conserved network of molecular interactions within the catalytic pocket (Figure 3B). Specifically, hydrogen bonds are formed with residues Ile392, Asn563, Ser525, and Asp409, corresponding to Ile378, Asn549, Ser511, and Asp395 in the human enzyme. These interactions anchor STM2457 within the active site. Additional stabilization arises from hydrophobic contacts and cation–π stacking interactions involving Arg550 (equivalent to Arg536 in human), a residue lining the SAM-binding cleft. Finally, structural superposition of the docked *A. aegypti* METTL3/METTL14–STM2457 complex with the human crystal structure revealed an almost perfect overlay of both the inhibitor and the catalytic residues (Figure 3C). The high degree of conservation in both amino acid residues and binding geometry strongly suggests that STM2457 functions as a cross-species inhibitor of METTL3.

**Figure 3.**
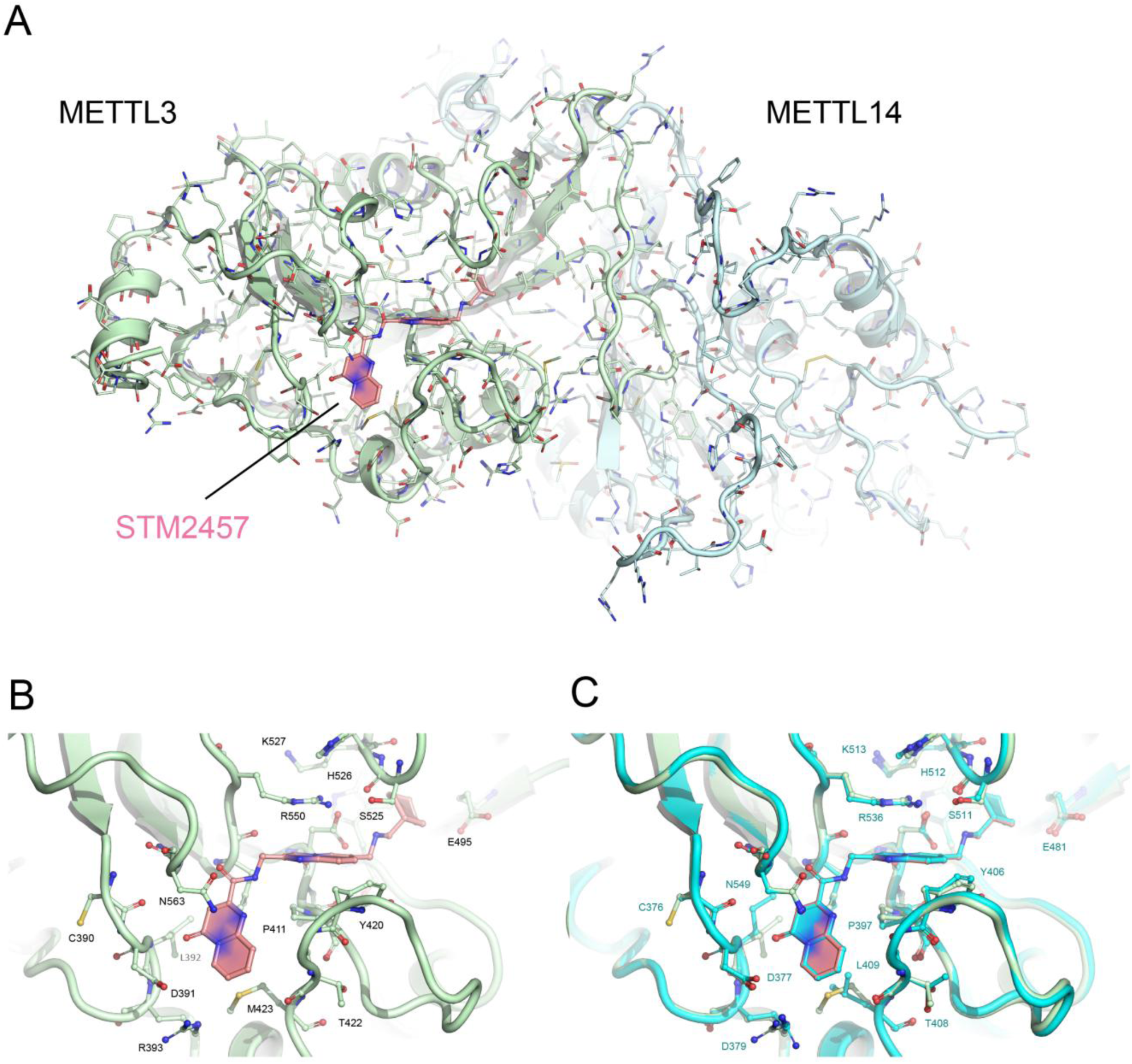
Docking of STM2457 to the modeled *A. albopictus* METTL3/METTL14 heterodimer, based on PDB 7O2I and generated via AlphaFold and AutoDock Vina. (A) Overall structural model of the dimeric *A. albopictus* METTL3 (green) and METTL14 (cyan) heterodimer, with STM2457 depicted as sticks in pink. (B) Close-up view of the METTL3/14 active site showing STM2457 nestled within the SAM-binding pocket. Key residues forming hydrogen bonds and hydrophobic contacts are labeled. (C) Superposition of the modeled *A. albopictus* complex with the human METTL3/METTL14·STM2457 crystal structure (PDB 7O2I, light blue). The near-perfect overlay of STM2457 and conserved active-site residues highlights the absolute conservation of binding interactions.

### Inhibition of m^6^A RNA methylation by STM2457 in C6/36 cells

To directly assess the impact of STM2457 on m^6^A RNA methylation in mosquito cells, C6/36 cells were treated with the inhibitor and total RNA was subsequently analyzed. Dot blot analyses revealed a consistent and significant reduction in global m^6^A levels in STM2457-treated cells compared to control conditions (Figure 4A and B), demonstrating effective inhibition of m^6^A RNA methylation. These results were independently validated by LC–MS analysis, which confirmed a significant decrease in m^6^A levels in total RNA isolated from STM2457-treated C6/36 cells (Figure 4C). Importantly, to evaluate the specificity of STM2457-mediated inhibition, the N^2,6^-dimethyladenosine (m^2,6^A) modification, predominantly found in ribosomal RNA, was simultaneously quantified by LC–MS. In contrast to m^6^A, m^2,6^A levels remained unchanged following STM2457 treatment (Figure 4D). These findings demonstrate that STM2457 effectively and specifically inhibits m^6^A RNA methylation in C6/36.

**Figure 4.**
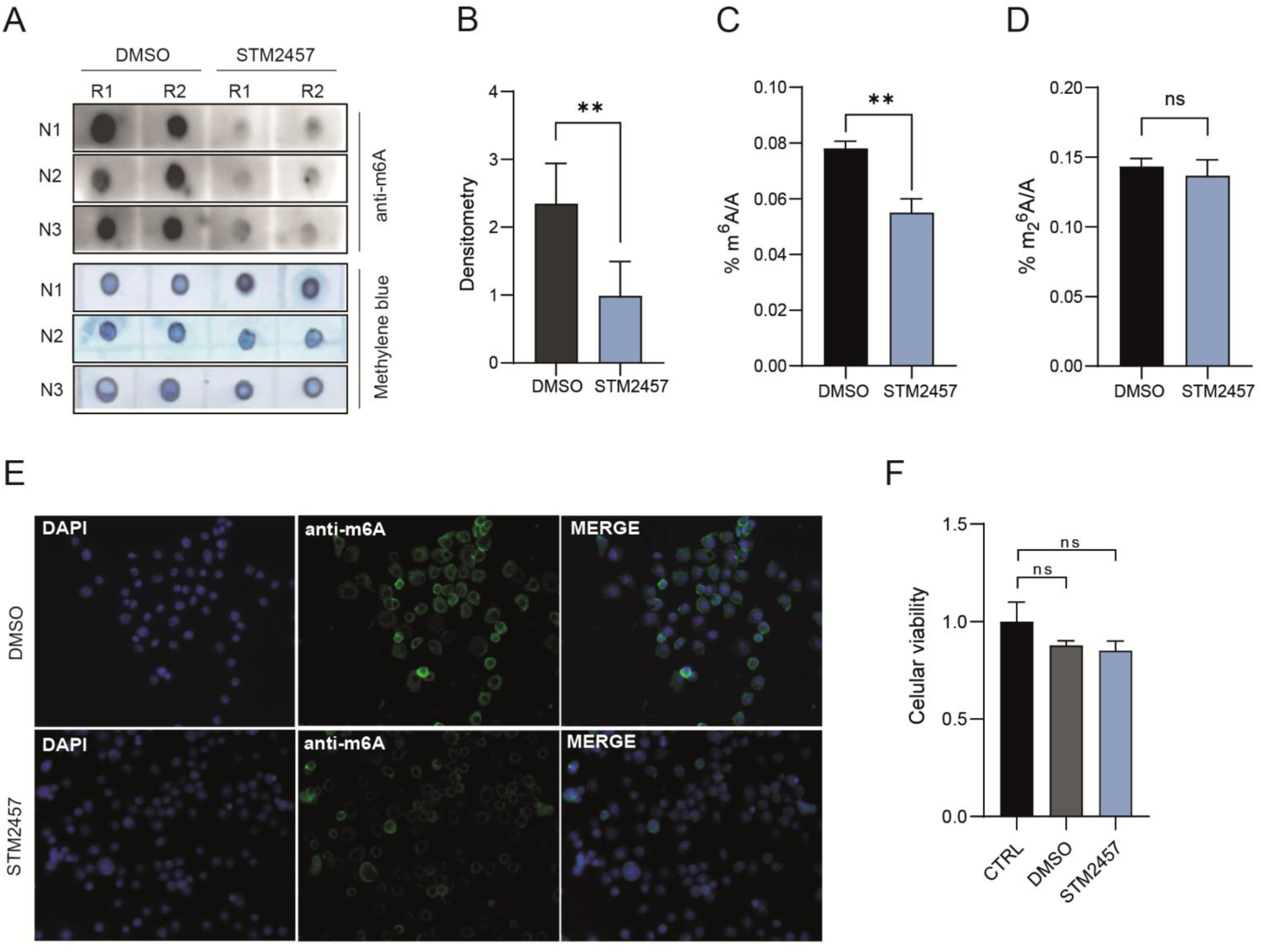
STM2457 selectively inhibits m^6^A RNA methylation in C6/36 cells. (A) Dot blot analysis of total RNA isolated from *A. albopictus* C6/36 cells treated with STM2457 at concentrations of 40 µM for 48 h. Membranes were probed with a monoclonal antibody specific for m^6^A to assess global m6A RNA levels (upper panel), followed by methylene blue staining to verify equal RNA loading (lower panel). N1-N3 refere to independent experimental replicates. R1 and R2 refere to two independente sample replicates. (B) Quantitative densitometric analysis of m^6^A dot blot signals shown in panel A. m^6^A signal intensities were normalized to methylene blue staining and expressed relative to DMSO-treated control cells. (C, D) Liquid chromatography–mass spectrometry (LC-MS) analysis of total RNA extracted from C6/36 cells treated with either DMSO or STM2457. (C) Quantification of m^6^A levels revealed a significant reduction in m^6^A abundance upon STM2457 treatment, expressed as percentage inhibition relative to controls. (D) In contrast, LC-MS analysis of m ^6^A showed no detectable change following STM2457 treatment, indicating specificity of STM2457 toward m^6^A methylation. (E) Immunofluorescence analysis of *A. albopictus* C6/36 cells treated with STM2457 (40 µM) for 48 h. Cells were fixed and stained with a monoclonal antibody specific for m^6^A to assess global m^6^A RNA levels at the single-cell level. Nuclei were counterstained with DAPI. Representative images are shown. (F) ATP-based cell viability assay of C6/36 cells treated with STM2457 (40 µM) for 48 h. Cellular ATP levels were quantified as a measure of metabolic activity and cell viability and are expressed relative to DMSO-treated controls. Statistical significance was determined using Student’s *t*-test

To further validate the inhibition of m^6^A RNA methylation by STM2457 in C6/36 cells, we performed immunofluorescence microscopy using anti-m^6^A antibodies. Consistent with the dot blot and LC–MS analyses, STM2457-treated cells exhibited a marked reduction in m^6^A-associated fluorescence intensity when compared to DMSO-treated control cells (Figure 4E), providing independent cellular-level confirmation of m^6^A inhibition.

Importantly, assessment of cell viability under the STM2457 concentrations used in these experiments revealed no detectable cytotoxic effects (Figure 4F), indicating that the observed reduction in m^6^A signal is not a consequence of compromised cell vitality.

### m^6^A inhibition leads to increased viral load in C6/36 cells

To investigate the functional impact of m^6^A RNA methylation on arbovirus replication in mosquito cells, C6/36 cells were treated with the METTL3 inhibitor STM2457 prior to infection with ZIKV. Notably, STM2457-treated cells exhibited markedly higher viral titers as determined by plaque assay for ZIKV (Figure 5A). In agreement with these results, quantification of viral RNA by qRT-PCR revealed a corresponding increase in viral genome levels following m^6^A inhibition (Figure 5B).

**Figure 5.**
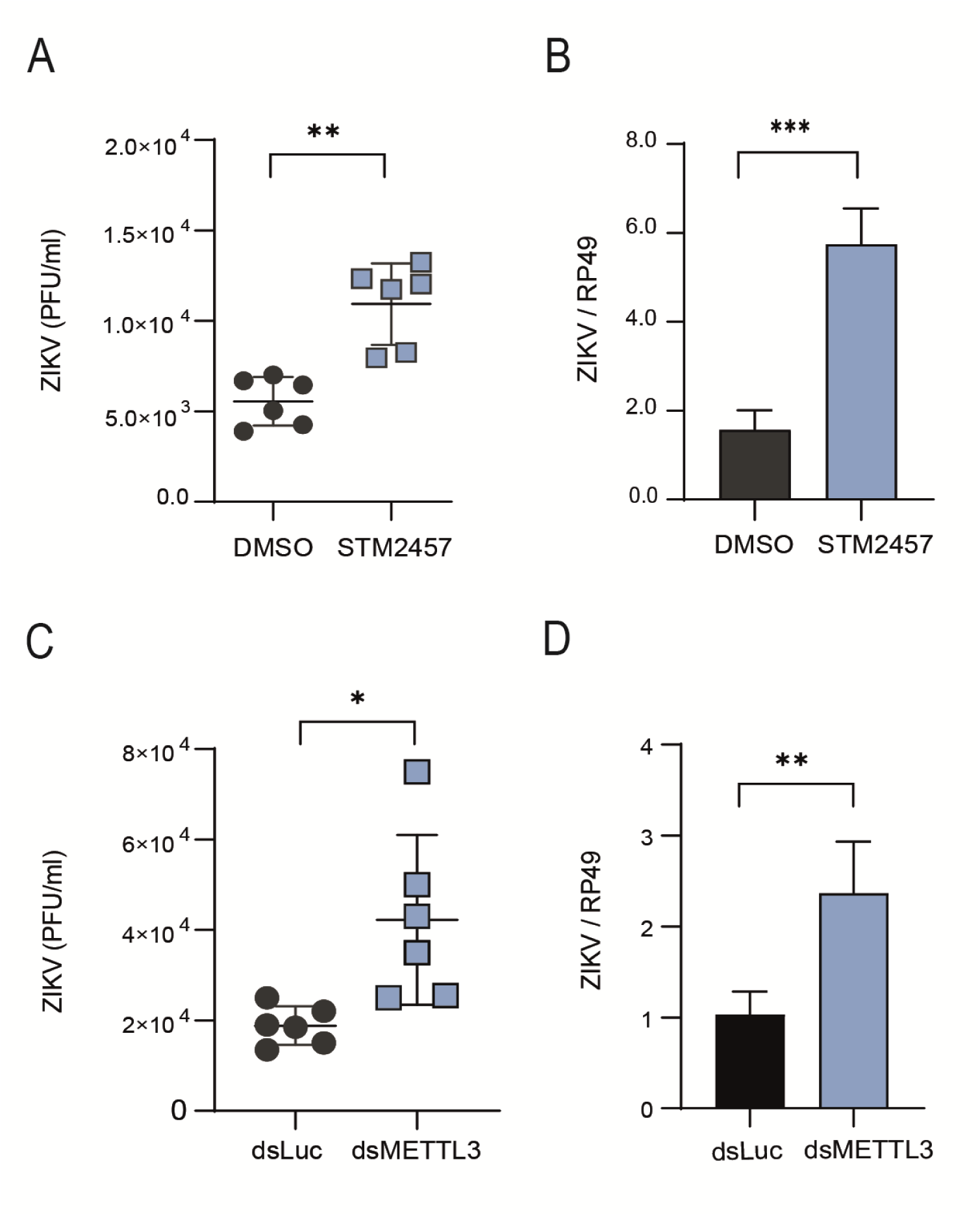
m^6^A RNA methylation acts as an antiviral mechanism in C6/36 cells. (A,B) *A. albopictus* C6/36 cells were pretreated with either dimethyl sulfoxide (DMSO) or the METTL3 inhibitor STM2457 (40 µM) for 48 hours prior to infection with ZIKV. Following pretreatment, cells were infected with ZIKV and maintained in culture for 3 days, after which the indicated analyses were performed. (C,D) A. *albopictus* C6/36 cells were transfected with double-stranded RNA (dsRNA) targeting METTL3 or with dsRNA corresponding to the irrelevant luciferase gene as a negative control. Viral replication was assessed by plaque assay (A and C) and quantitative RT-PCR (B and D). Data are presented as mean ± SEM, and statistical significance was determined using Student’s *t*-test. Genetic silencing of AaMETTL3 phenocopied the effects observed upon pharmacological inhibition of m^6^A RNA methylation, supporting an antiviral role for METTL3-mediated m^6^A modification.

For further confirmation, we used dsRNA-mediated silencing of *METTL3* in *A. aegypti* Aag-2 cells. This resulted in viral loads that were indistinguishable from those observed following treatment with the m^6^A inhibitor STM2457, further confirming that the increase in viral load is specifically associated with inhibition of the m^6^A methylation pathway (Figure 5C and D). Together, these findings demonstrate that inhibition of m^6^A RNA methylation enhances arbovirus replication in *Aedes* mosquitoes, supporting a role for m^6^A-dependent post-transcriptional regulation as a restrictive host mechanism during ZIKV infection in mosquito cells.

### Transcriptomic profiling of *A. albopictus* C6/36 cells upon STM2457 treatment

To further investigate the effects of STM2457 treatment, we used RNA sequencing (Figure 6). This generated a high-quality and robust dataset from both experimental conditions, with an average of 21.36 million reads obtained from DMSO-treated control samples and 22.56 million reads from STM2457-treated samples (https://www.ncbi.nlm.nih.gov/sra/PRJNA1417765). Read alignment to the *A. albopictus* reference genome showed similarly high mapping efficiencies between groups, with 89.6 % of reads (approximately 19.15 million) mapping in the DMSO condition and 89.2 % of reads (approximately 20.14 million) mapping in the STM2457-treated condition, indicating reliable sequencing depth and alignment quality.

**Figure 6.**
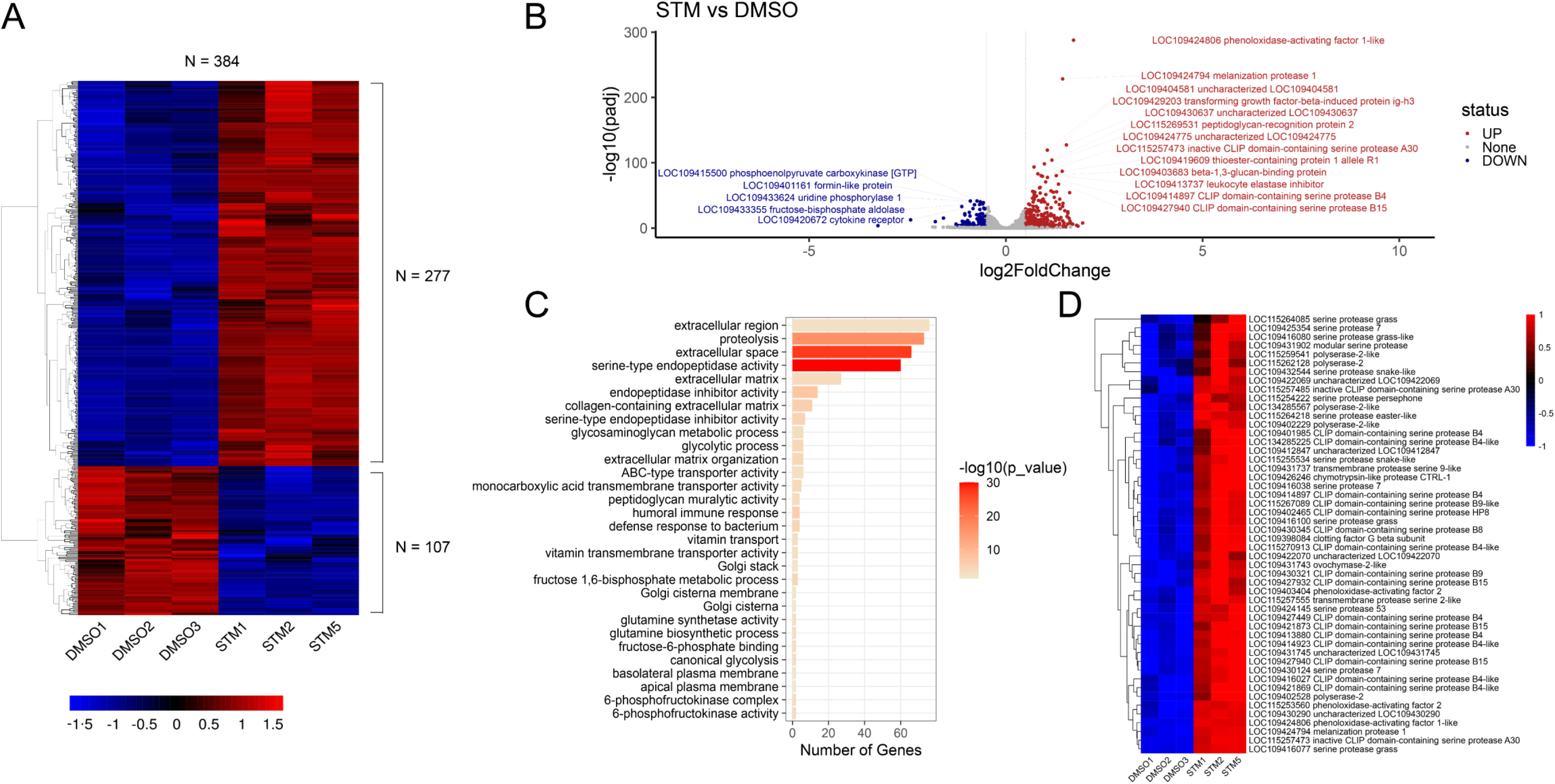
Transcriptomic profiling of *A. albopictus* C6/36 cells upon treatment with STM2457. (A) Heatmap of the 384 differentially expressed genes (DEGs) identified between STM2457-treated and DMSO control cells, comprising 277 up-regulated and 107 down-regulated transcripts. Each row represents a single gene, and expression values are scaled as z-scores to highlight relative differences across samples. (B). Volcano plot depicting the distribution of DEGs. Significantly up-regulated genes are shown in red, down-regulated genes in blue, and non-significant genes in grey. Dashed horizontal line indicates the significance threshold (p-value < 0.01), while vertical dashed lines mark the log₂ fold-change cutoffs (–0.5 and +0.5). Selected top gene IDs and their annotations are displayed to highlight biologically relevant changes. (C). Gene Ontology (GO) enrichment analysis of differentially expressed genes. The X-axis shows the number of DEGs associated with each enriched GO term, while the color gradient represents statistical significance (–log₁₀ p-value). Enriched terms include immune- and protease-related functions. (D) Heatmap of the subset of 60 DEGs assigned to the enriched GO category “serine-type endopeptidase activity.” Expression values are presented as z-scores for each gene, illustrating clear transcriptional modulation of serine protease–related genes in response to STM2457 treatment.

Differential expression analysis identified a total of 384 genes significantly modulated upon STM2457 treatment compared to DMSO controls. Of these, 277 genes were up-regulated and 107 genes were down-regulated (Figure 6A). Notably, a substantial fraction of the up-regulated genes was associated with components of the insect innate immune system. Among these, we observed increased expression of genes encoding key regulators of melanization and proteolytic cascades, including LOC109424806 (phenoloxidase-activating factor 1–like), LOC109424794 (melanization protease 1), as well as multiple CLIP-domain-containing serine proteases, such as LOC115257473 (inactive CLIP domain-containing serine protease A30), LOC109414897 (CLIP domain-containing serine protease B4), and LOC109427940 (CLIP domain-containing serine protease B15) (Figure 6B). The induction of these genes suggests a coordinated activation of protease-driven pathways typically involved in melanization and other immune-related defense responses.

Consistent with these observations, Gene Ontology (GO) enrichment analysi**s** revealed a significant overrepresentation of immune-associated functional categories among the differentially expressed genes. One of the most strongly enriched GO terms was “serine-type endopeptidase activity” (p < 0.01), which encompassed 60 differentially expressed genes (Figure 6C and D). This pronounced enrichment underscores the prominence of serine proteases within the transcriptional response elicited by STM2457 treatment in C6/36 cells.

### m^6^A methylomes of C6/36 cells and arboviruses

To map m^6^A RNA within host transcripts and to assess whether arboviral genomes are m^6^A-modified, we performed GLORI (Global m^6^A Level and Isoform-specific) [47] analyses on RNA derived from C6/36 cells infected with ZIKV or CHIKV. This approach enabled transcriptome-wide absolute quantification of m^6^A levels at single-base resolution. Data analysis revealed only background levels of non-conversion in the ZIKV virus RNAs (Figure 7A). Similar results were obtained from parallel analyses of CHIKV-infected C6/36 cells (Supplementary Figure 4), strongly suggesting that these arboviral RNAs are not m^6^A-modified in C6/36 cells. In contrast, we detected 11.569 m^6^A sites in ZIKV-infected C636 cells, and 10.559 sites in CHIKV-infected C636 cells, which were highly enriched within the canonical DRACH consensus motif (Figure 7B, Supplementary Figure 4C). Within the 10 most frequent pentanucleotide motifs, canonical DRAC(H) motifs were strongly overrepresented (Figure 7C, Supplementary Figure 4C). Also, higher and more evenly distributed m^6^A methylation levels were detected in the DRAC(H) motifs, while non-canonical motifs were lowly methylated (Figure 7D, Supplementary Figure 4D). Metagene plots showed that the majority of m^6^A sites were in the CDS, with a prominent enrichment around the 5’-end (Figure 7E, Supplementary Figure 4E).

**Figure 7.**
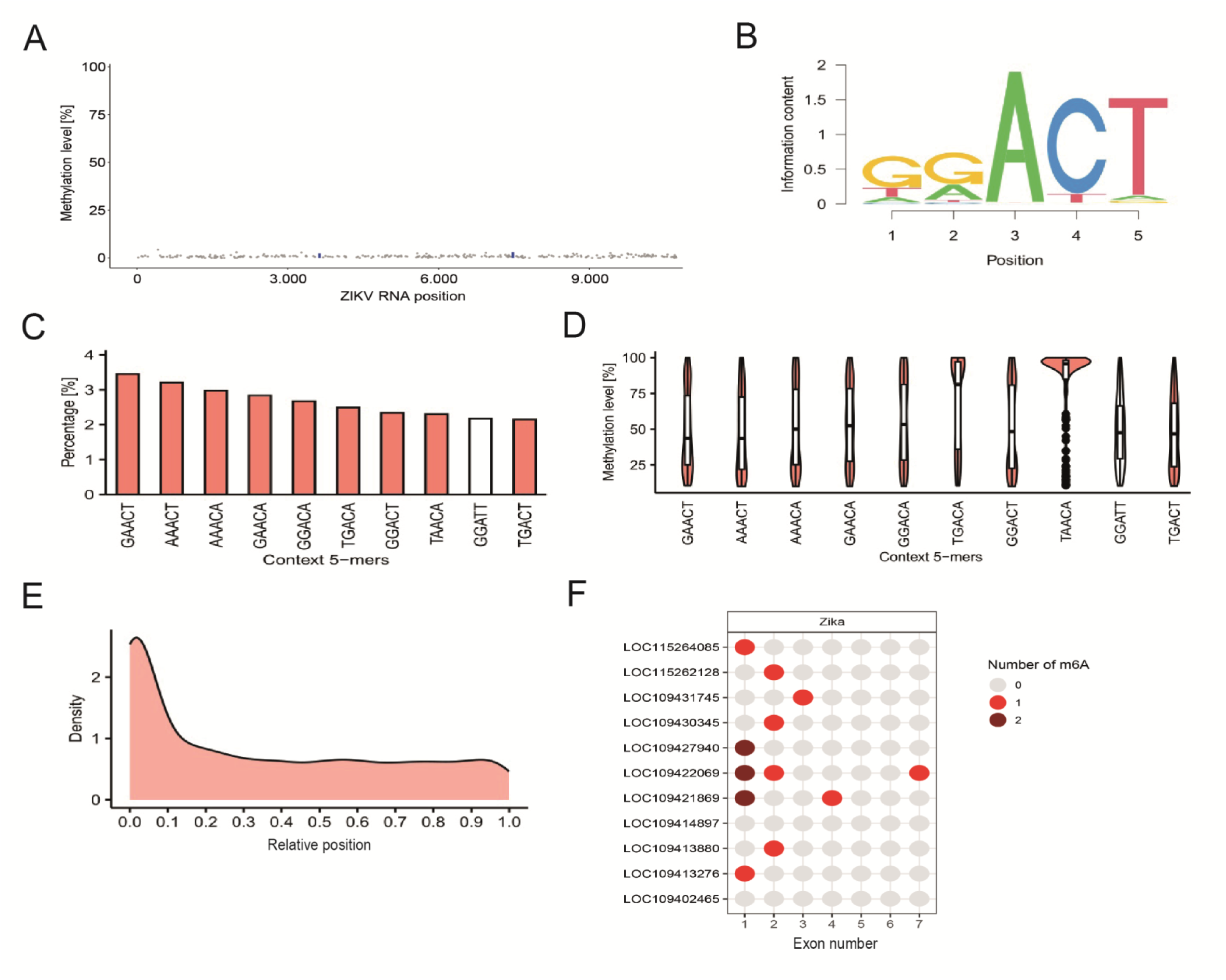
The m^6^A epitranscriptomic landscape of ZIKV-infected C6/36 cells. (A) GLORI-seq analysis of ZIKV viral RNAs. Grey dots indicate non-conversion levels of adenines in DRAC sites. Blue bars indicate potential methylation sites with a ratio >0.01 and Padj <0.05. Note that only a small fraction of sites exceeds these cutoffs and that their ratios are consistently low. (B) Sequence motif analyses using detected m^6^A sites in virus-infected C636 cells. m^6^A sites were primarily detected in the DRAC(H) consensus sequence motif. (C) Frequencies of the 10 most-detected motifs in virus-infected C636 cells. (D) Quantification of the methylation level in the 10 most-detected motifs. (E) Metagene plot depicting the positional distribution of detected m⁶A sites along normalized transcripts. m⁶A modifications are predominantly enriched within the CDS, with a higher density toward the 5′ region of the coding sequence. (F) Dotplot displaying distribution and number of detected m^6^A methylation sites in serine protease transcripts that were upregulated following pharmacological inhibition of METTL3 with STM2457, as identified by RNA-Seq. The results reveal that m^6^A modifications are broadly distributed throughout the CDS, without strong confinement to transcript termini, indicating preferential deposition within protein-coding regions under the analyzed conditions.

To further integrate the GLORI and RNA-seq datasets, we examined the presence of m^6^A modification sites within transcripts encoding selected domain-containing serine proteases that were transcriptionally modulated upon STM2457 treatment. The dotplots showed that the majority of m^6^A sites were in the CDS, with a prominent enrichment around the 5’-end (Figure 7F, Supplementary Figure 4F).

### Covalent docking of AEBSF to *A. aegypti* serine proteases

AEBSF is a water-soluble, irreversible serine protease inhibitor that covalently modifies the catalytic serine residue [48]. To evaluate its inhibitory potential against mosquito CLIP-domain serine proteases (CLIPs), we performed covalent docking simulations using selected *A. aegypti* CLIPs (B4, B9, B15, Sp7, Sp-g, and Sp-s; Figure 8A). Docking predicted stable covalent adduct formation between AEBSF and the catalytic serine in all CLIPs except Sp7, in which the catalytic serine is replaced by glycine, precluding covalent interaction. In catalytically competent CLIPs, AEBSF adopted a conserved orientation that positioned its reactive sulfonyl group near the nucleophilic serine, supporting irreversible inhibition.

**Figure 8.**
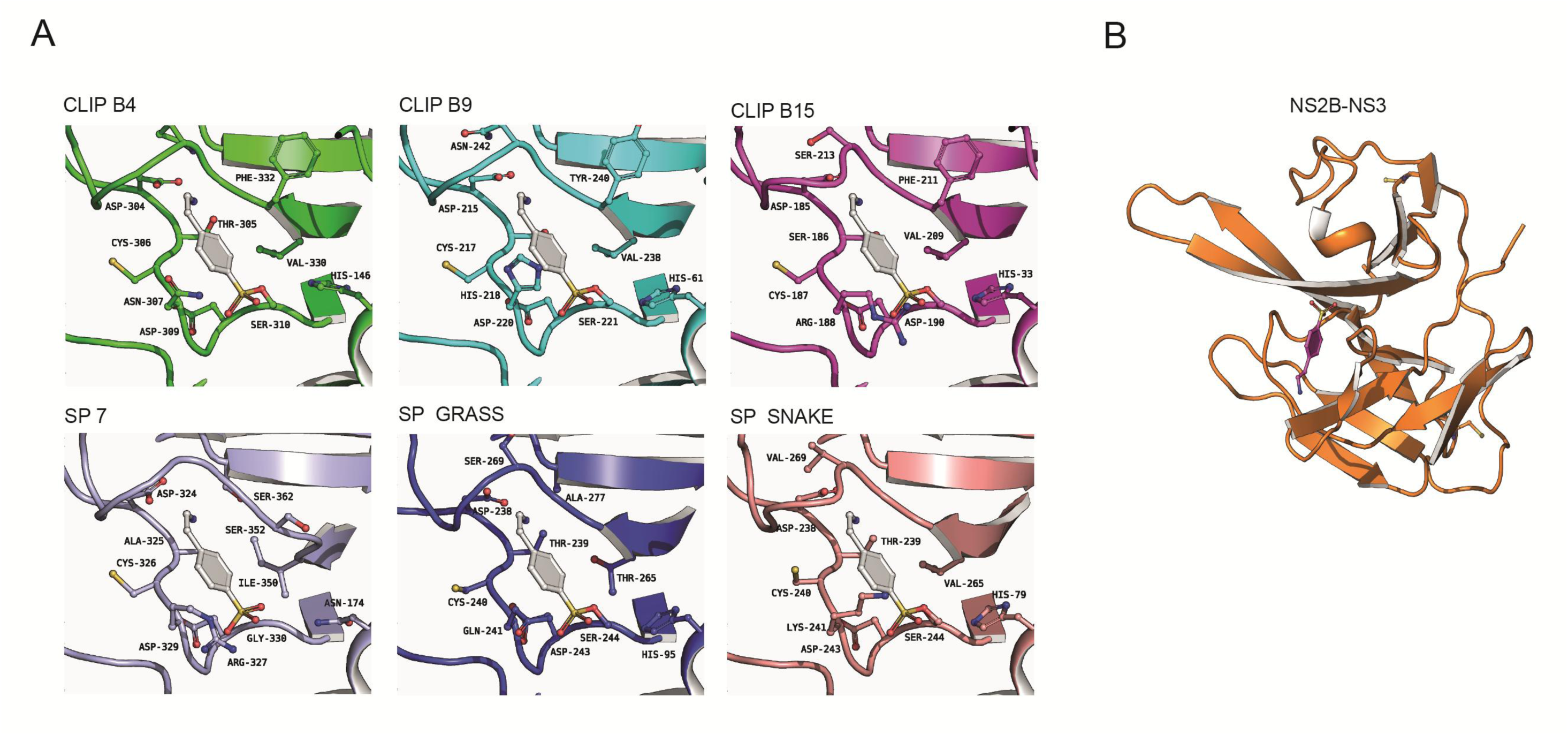
Docking of AEBSF into the 3D models of host and virus proteases. (A) Predicted binding poses of AEBSF (4-(2-Aminoethyl)benzenesulfonyl fluoride) within the catalytic sites of the CLIP serine proteases B4, B9, B15, Sp7, Sp-g, and Sp- s. Each panel shows the best docking conformation obtained from molecular docking simulations, highlighting the spatial orientation of the inhibitors relative to the catalytic triad residues. The inhibitor is depicted in sticks representation, while the proteases are shown as cartoon ribbons. Key residues stabilizing the inhibitor are indicated and labeled. The dashed yellow line highlights salt-bridge interactions. Statistical significance was determined using Student’s *t*-test. (B) Predicted binding poses of AEBSF (4-(2-Aminoethyl)benzenesulfonyl fluoride) within the catalytic sites of the NS2B-NS3 ZIKV protease. The best docking conformation obtained from molecular docking simulations is reported, highlighting the spatial orientation of the inhibitor relative to the catalytic triad residues. The inhibitor is depicted in sticks representation, while the protease is shown as cartoon ribbons. Key residues stabilizing the inhibitor are indicated and labeled. The dashed yellow line highlights salt-bridge interactions.

We next assessed potential interactions with the ZIKV NS2B–NS3 protease complex (Figure 8B). In contrast to mosquito CLIPs, AEBSF showed lower predicted affinity and less stable positioning within the viral active site, with docking poses that did not favor efficient covalent engagement. Structural analysis indicated that the NS2B–NS3 binding pocket is less compatible with productive AEBSF stabilization, particularly regarding orientation of the inhibitor’s amino group. Accordingly, predicted interactions with the viral protease were weaker than with host CLIPs.

Importantly, AEBSF treatment up to 400 µM did not affect C6/36 cell viability (Supplementary Figure 3), indicating that subsequent effects are not attributable to cytotoxicity.

### Intracellular serine proteases modulate ZIKV replication in C6/36 cells

Serine proteases can act as proviral factors at multiple stages of infection. In vertebrate systems, extracellular or plasma membrane–associated serine proteases enhance viral attachment and entry [49–52]. To determine whether a similar mechanism operates in mosquito cells, C6/36 cells were pretreated with AEBSF prior to ZIKV exposure, and viral binding and internalization were quantified. As shown in Figure 9A, serine protease inhibition did not significantly affect viral attachment or entry, indicating that extracellular or membrane-associated serine proteases are not required for ZIKV entry into mosquito cells.

**Figure 9.**
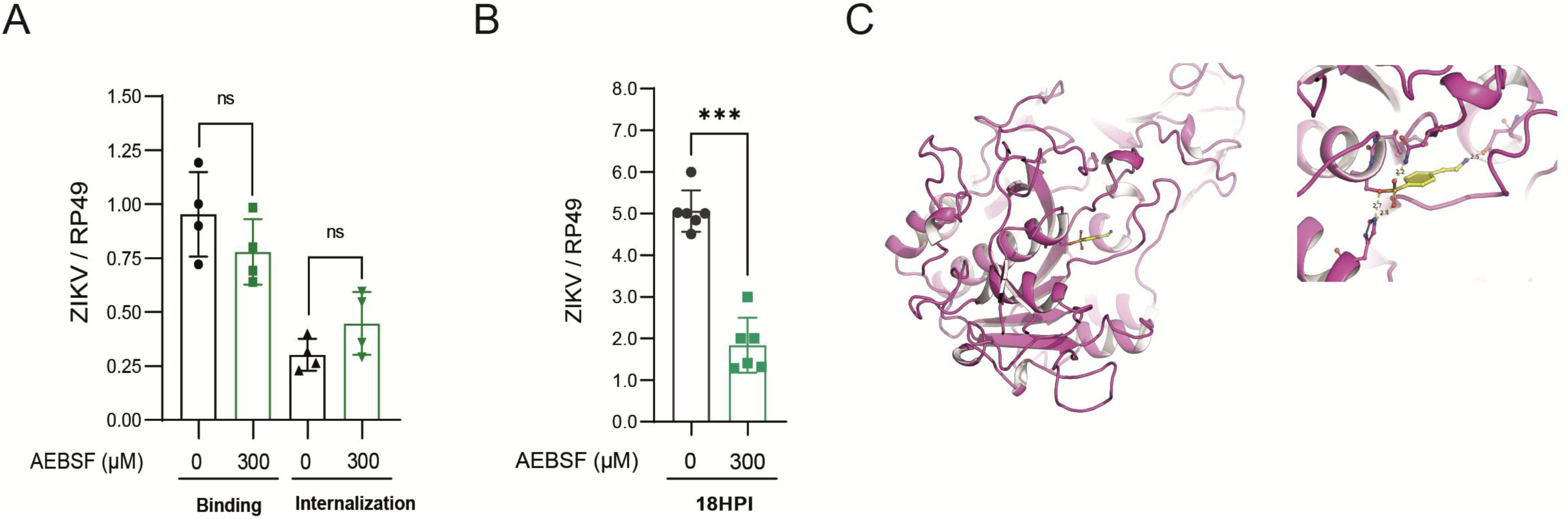
Effects of AEBSF-mediated inhibition on host and viral serine proteases during ZIKV infection. (A,B) Quantitative PCR (qPCR) analysis of ZIKV RNA levels in *A. albopictus* C6/36 cells under conditions of serine protease inhibition. Cells were infected with ZIKV and subsequently treated with the serine protease inhibitor AEBSF (300 µM), which was maintained in the culture medium for 18 hours post-infection (A). In parallel, cells were pretreated with AEBSF (300 µM) for 24 hours prior to ZIKV infection (B). Viral RNA levels were quantified by qPCR and normalized to control conditions. Data are presented as relative viral RNA abundance. Statistical significance was determined using Student’s *t*-test. (C,D) Predicted binding poses of AEBSF (4-(2-Aminoethyl)benzenesulfonyl fluoride) within the catalytic sites of *A. aegypti* Furin protease domain (residues 271–743). The best docking conformation obtained from molecular docking simulations is reported, highlighting the spatial orientation of the inhibitors relative to the catalytic triad residues. The inhibitor is depicted in sticks representation, while the protease is shown as cartoon ribbons. Key residues stabilizing the inhibitor are indicated and labeled. The dashed yellow line highlights salt-bridge interactions.

We next evaluated post-entry stages. Infected C6/36 cells were treated with AEBSF for 18 hours, and viral replication was quantified (Figure 9B). Under these conditions, serine protease inhibition reduced viral load by more than 50% compared to untreated controls, demonstrating that serine proteases contribute to ZIKV replication after entry.

Given the pleiotropic activity of AEBSF, we investigated furin as a potential intracellular target. Covalent docking using an AlphaFold3-predicted structural model of the *A. aegypti* furin protease domain (residues 271–743) showed that AEBSF adopts a productive binding pose within the active site, positioning its sulfonyl fluoride group near the catalytic serine (Figure 9C). This configuration supports covalent enzyme–inhibitor complex formation, consistent with direct inhibition of mosquito furin.

## Discussion

In this study, we identify m^6^A RNA methylation as a key epitranscriptomic regulator of arbovirus infection in *A. albopictus* and establish serine proteases as a mechanistic link between m^6^A regulation and viral replication.

We show that *A. albopictus* encodes a complete and highly conserved m^6^A machinery, which is consistent with previous reports identifying functional m^6^A pathways in insects and other invertebrates, supporting the notion that m^6^A-mediated RNA regulation is an evolutionarily ancient mechanism [53]. Despite this conservation, we observed no detectable modulation of the m^6^A machinery following infection with ZIKV in mosquito cells. This contrasts with mammalian systems, where viral infection can induce dynamic changes in m^6^A levels or in the expression of m^6^A writers, erasers, or readers [1–9]. Instead, our data support a model in which m^6^A operates as a basal regulatory mechanism controlling RNA fate, in agreement with foundational studies defining m^6^A as a widespread modulator of mRNA stability and translation [9,54,55].

Using both pharmacological and genetic approaches, we demonstrate that inhibition of m^6^A RNA methylation significantly enhances ZIKV replication in mosquito cells revealing that m^6^A exerts a net antiviral effect in *A. albopictus* cells. Antiviral roles for m^6^A have been reported for several RNA viruses, including flaviviruses and alphaviruses, although the directionality of m^6^A effects is context dependent [2,3]. In insects, disruption of m^6^A has been shown to increase susceptibility to viral infection, supporting a role for epitranscriptomic regulation in vector competence [56–58].

Transcriptomic profiling revealed that inhibition of m^6^A results in strong upregulation of serine proteases, including CLIP-domain proteases. RNA methylome analysis using single-nucleotide–resolution GLORI further identified m^6^A modifications within multiple serine protease transcripts, strongly suggesting that these genes are direct m^6^A targets. The m^6^A modification is known to regulate mRNA decay and translation via YTH-domain reader proteins, thereby fine-tuning gene expression programs involved in stress responses and immunity [59]. Our data extend this regulatory paradigm to mosquito serine proteases.

Our GLORI analyses further revealed no detectable m^6^A modification within the genomes of ZIKV or CHIKV in mosquito cells. This observation contrasts with earlier reports from mammalian systems, in which flaviviral and alphaviral RNAs were shown to harbor m^6^A sites implicated in the regulation of viral gene expression, replication, and immune recognition [20–22,56,60]. Our findings suggest that, in mosquito cells, arboviral RNAs either evade m^6^A deposition or that any such modifications occur at levels below the detection threshold under the experimental conditions tested. Importantly, our GLORI-based results are independently supported by a recent comprehensive study employing multiple antibody-independent approaches, which likewise failed to detect m^6^A modifications in CHIKV or dengue virus (DENV) transcripts [24].

CLIP serine proteases are central components of insect immune signaling, particularly melanization cascades and immune amplification pathways [61,62]. However, their activity must be tightly controlled, as excessive protease activation can disrupt immune homeostasis and cause collateral cellular damage [63–66]. We therefore propose that m^6^A normally constrains serine protease expression to prevent maladaptive proteolytic activity.

Consistent with this model, dysregulated serine protease expression correlated with enhanced viral replication. Although serine proteases are classically associated with host defense, numerous viruses exploit host proteolytic environments to promote entry, replication, or maturation [49,67,68]. Excessive protease activity may also degrade immune signaling components, thereby indirectly promoting viral replication [69,70].

An important finding of this study is the impact of serine protease inhibition on ZIKV infection in mosquito cells. Pharmacological treatment with the irreversible serine protease inhibitor AEBSF resulted in a significant reduction in ZIKV replication, indicating that serine protease activity contributes to efficient viral infection. Molecular docking analyses provide mechanistic insight into this observation, revealing that AEBSF binds with high affinity to multiple mosquito serine proteases, while exhibiting comparatively weaker predicted interactions with the ZIKV NS2B–NS3 protease. The viral NS2B–NS3 protease is essential for flaviviral polyprotein processing and represents a well-established antiviral target [71–73]. Although AEBSF is predicted to interact less efficiently with the viral protease than with host enzymes, its partial engagement with NS2B–NS3 raises the possibility that AEBSF may exert a modest direct antiviral effect in addition to its primary action on host proteases.

Further analysis indicated a stage-specific role for serine proteases during ZIKV infection of mosquito cells. Unlike vertebrate systems, where extracellular or plasma membrane–associated serine proteases can facilitate viral entry [52,74], inhibition of serine protease activity in C6/36 cells did not affect viral binding or internalization, indicating that these enzymes are not involved in ZIKV entry in mosquitoes. In contrast, inhibition of serine proteases after viral entry significantly reduced viral replication, demonstrating that intracellular serine proteases act as proviral factors at post-entry stages of infection. These results highlight host-specific differences in the role of serine proteases during ZIKV infection while supporting a conserved requirement for intracellular serine protease activity in efficient viral replication.

Another important consideration in the interpretation of these results relates to the specificity of the serine protease inhibitor AEBSF. Although AEBSF is expected to efficiently inhibit CLIP family proteases in insects, its broad reactivity and lack of strict specificity raise the possibility that additional host serine proteases, such as furin, may also be affected. Furin-mediated cleavage is essential for ZIKV maturation within the trans-Golgi network in mammalian cells [75]. Consistent with this notion, our *in silico* covalent docking analysis revealed that AEBSF adopts a productive binding pose within the active site of furin, positioning its reactive moiety in close proximity to the catalytic serine residue. This finding suggests that furin or furin-like proprotein convertases could represent additional intracellular targets of AEBSF, potentially contributing to the observed reduction in ZIKV infection. While these results do not establish direct inhibition of furin activity *in vivo*, they underscore the need to consider off-target effects of AEBSF when interpreting the antiviral phenotype and highlight the importance of future studies employing more selective genetic or pharmacological approaches to dissect the specific contributions of individual serine proteases.

These findings integrate with our epitranscriptomic data to support a broader model in which m⁶A RNA methylation functions as a constitutive regulator of mosquito permissiveness to arbovirus infection. In this model, depicted in Figure 10, m^6^A-mediated control restrains the expression of specific serine proteases, including CLIP-domain proteases, thereby limiting protease-dependent processes that favor viral replication. Disruption of m⁶A deposition, through inhibition of the METTL3 methyltransferase, leads to derepression of serine protease transcripts and creates a cellular environment more conducive to arbovirus infection.

**Figure 10.**
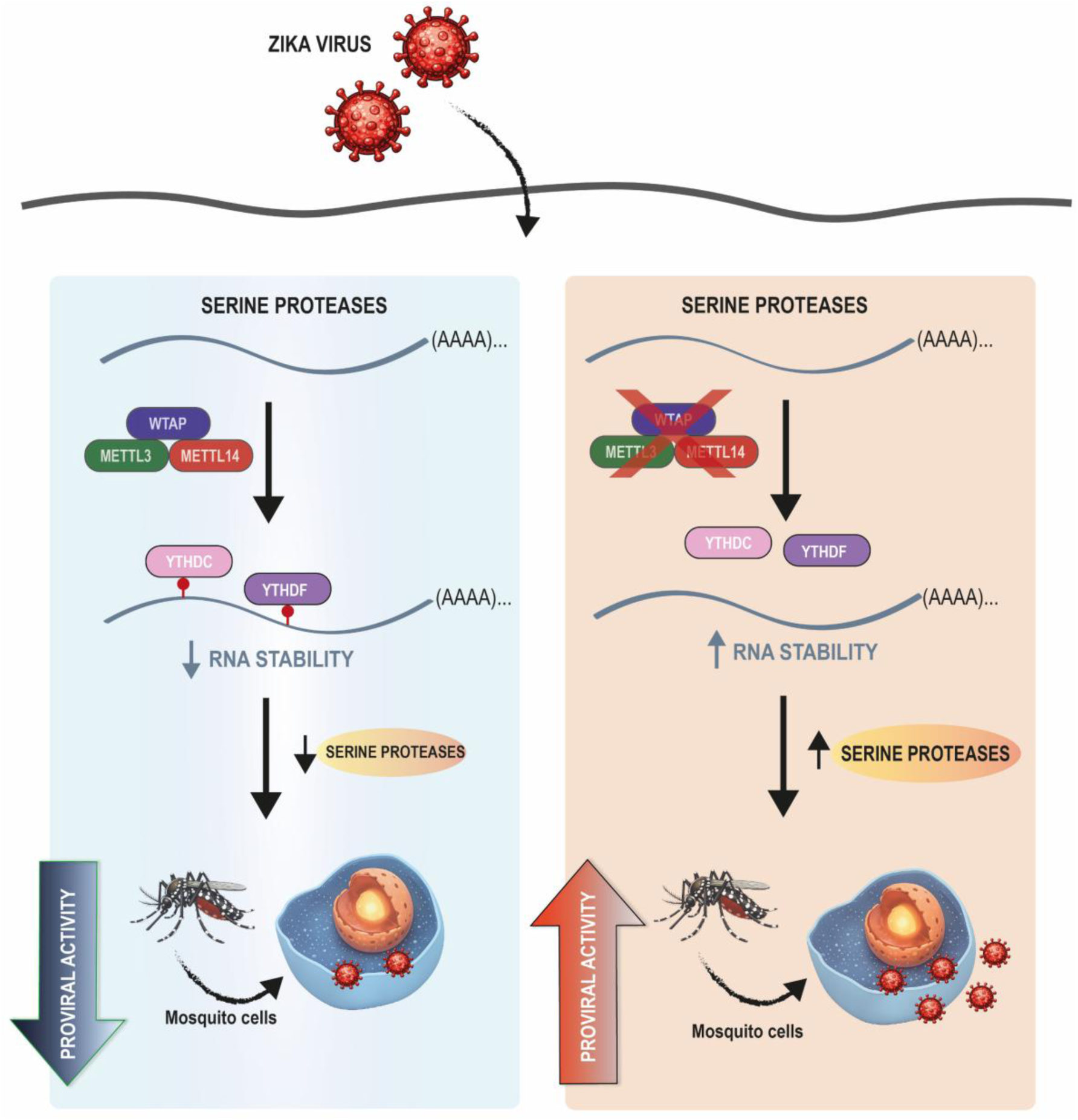
Proposed model for the role of m^6^A RNA methylation in the regulation of arbovirus infection in *Aedes* mosquitoes. Overall, our results indicate that serine proteases from *A. albopictus* act as proviral factors that support ZIKV infection, particularly at post-entry stages of the viral life cycle. Left panel: Under basal conditions, m^6^A RNA methylation limits the expression of serine proteases (CLIPs included) mRNAs, thereby maintaining the cellular environment of C6/36 cells in a state that is less permissive to ZIKV infection. Right panel: Reduction of m^6^A RNA methylation leads to increased stability and/or enhanced translation of serine protease transcripts. The resulting increase in serine protease expression and proteolytic activity restricts ZIKV infection.

At this stage, the precise molecular pathways through which serine protease inhibition limits ZIKV infection remain to be defined, including whether this effect reflects disruption of host protease–dependent processes, interference with viral protein maturation, or a combination of both. Thus, although our data establish a functional connection between m^6^A regulation, serine protease activity, and ZIKV infection, they also underscore a critical gap in our understanding of how proteolytic networks shape arbovirus–mosquito interactions. Future studies aimed at identifying the specific proteases and substrates involved will be essential to resolve the mechanistic basis of this regulation and to assess its relevance for vector competence and viral transmission *in vivo*.

## Acknowledgments

We thank Jaciara Miranda Freire for her invaluable support in maintaining the insectary and for providing outstanding technical assistance throughout the study. We also thank Dr. Martin Hengesbach and Dr. Martina Krämer (Institute of Pharmaceutical and Biomedical Sciences Johannes Gutenberg-University Mainz, Germany) for their support with the LC-MS analysis. This work utilized the computational resources of the NIH HPC Biowulf cluster (http://hpc.nih.gov). The contributions of the NIH author(s) were made as part of their official duties as NIH federal employees, are in compliance with agency policy requirements, and are considered Works of the United States Government. However, the findings and conclusions presented in this paper are those of the author(s) and do not necessarily reflect the views of the NIH or the U.S. Department of Health and Human Services.

## Author contributions

Conceptualization: AMA, MRF; Methodology: AMA, LT, JK, FB, SR, TB, LAT, LMK; Formal analysis: AMA, LT, JK, FB, SR, AP, DR, TB, LAT, JDPLY, VCC, FL, MRF; Investigation: AMA, LT, JK, FB, SR, AP, DR, TB, LAT, JDPLY, VCC; Data Curation: AMA, LT, JK, FB, SR, AP, DR, TB, LAT, JDPLY, VCC, APANE, KL, MRF; Writing original draft: AMA, LT, SR, AP, TB, FL, MRF; Writing review: AMA, LT, SR, AP, TB, FL, MRF; Supervision: MRF, LT, AP, DR, FL, MRF; Funding Acquisition: LT, AP, DR, FL, MRF; Project Administration: MRF.

## Data availability

The transcriptome data was deposited to the National Center for Biotechnology Information (NCBI) under BioProject PRJNA1417765 and Biosamples SAMN55014584 - SAMN55014589. The raw reads were deposited to the Short Reads Archive of the NCBI under accessions SRRSRR37075377 - SRRSRR37075382.

---

**Supplementary Figure 1.**
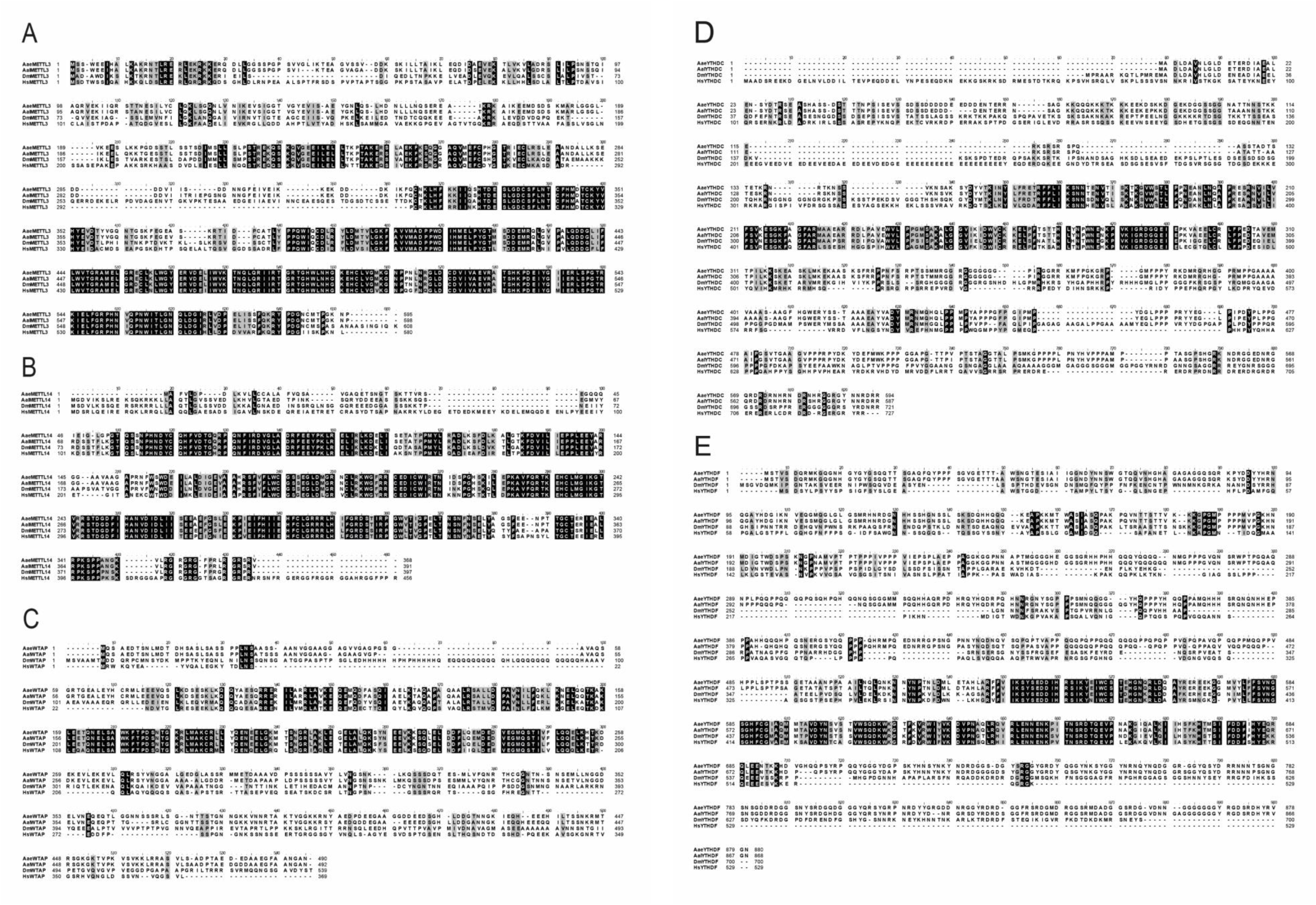
Multiple sequence alignment of the full-length METTL3, METTL14, WTAP, YTHDC, and YTHDF proteins from *A. aegypti*, *A. albopictus*, *Drosophila melanogaster*, and human. Amino acid residues shaded in gray indicate conserved positions among species, whereas residues shaded in black denote positions that are identical across all aligned sequences. This alignment highlights the high degree of evolutionary conservation of core m⁶A-associated factors across insects and vertebrates.

**Supplementary Figure 2.**
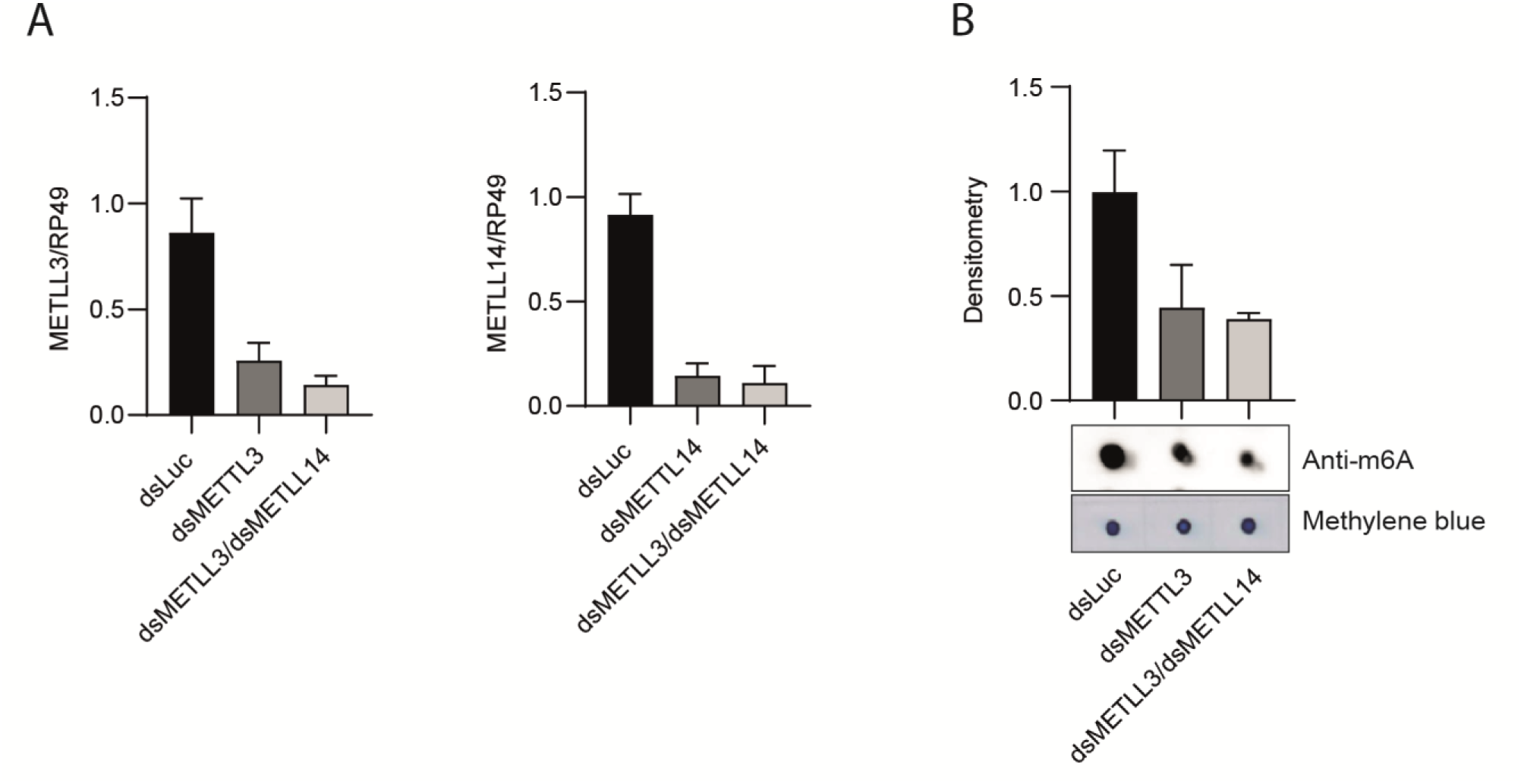
AaMETTL3 knockdown enhances arbovirus replication in Aag-2 cells. (A–F) *A. aegypti* Aag-2 cells were transfected with double-stranded RNA (dsRNA) targeting METTL3 (dsMETTL3) or a control luciferase dsRNA (dsLuc) and subsequently infected with ZIKV. dsRNA treatment was performed for 4 days prior to viral infection. Knockdown efficiency was evaluated by assessing AaMETTL3 transcript levels by RT-qPCR (A,B) and by measuring METTL3/METTL14 methyltransferase activity (C). Data are presented as mean ± SEM, and statistical significance was determined using Student’s *t*-test.

**Supplementary Figure 3.**
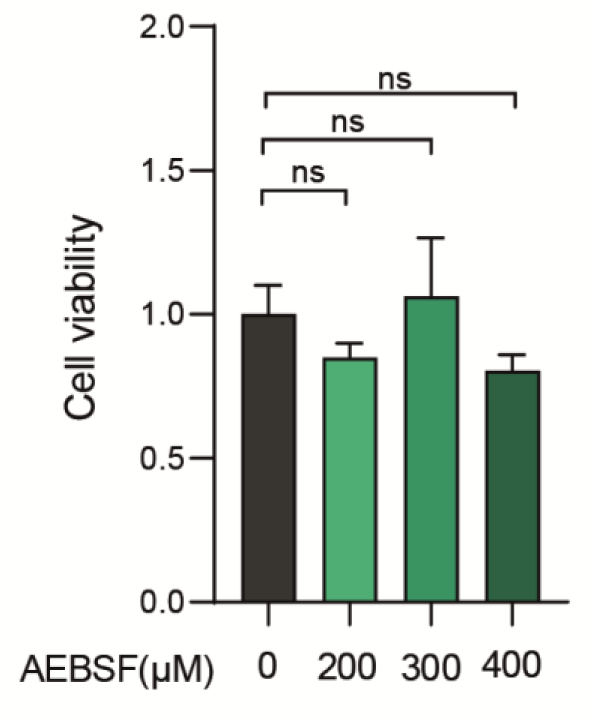
ATP-based cell viability assay of *A. albopictus* C6/36 cells treated with increasing concentrations of the serine protease inhibitor AEBSF for 48 h. Cellular ATP levels were measured as an indicator of metabolic activity and cell viability and are expressed relative to DMSO-treated control cells. No statistically significant differences in cell viability were observed across the tested AEBSF concentrations, as determined by Student’s *t*-test.

**Supplementary Figure 4.**
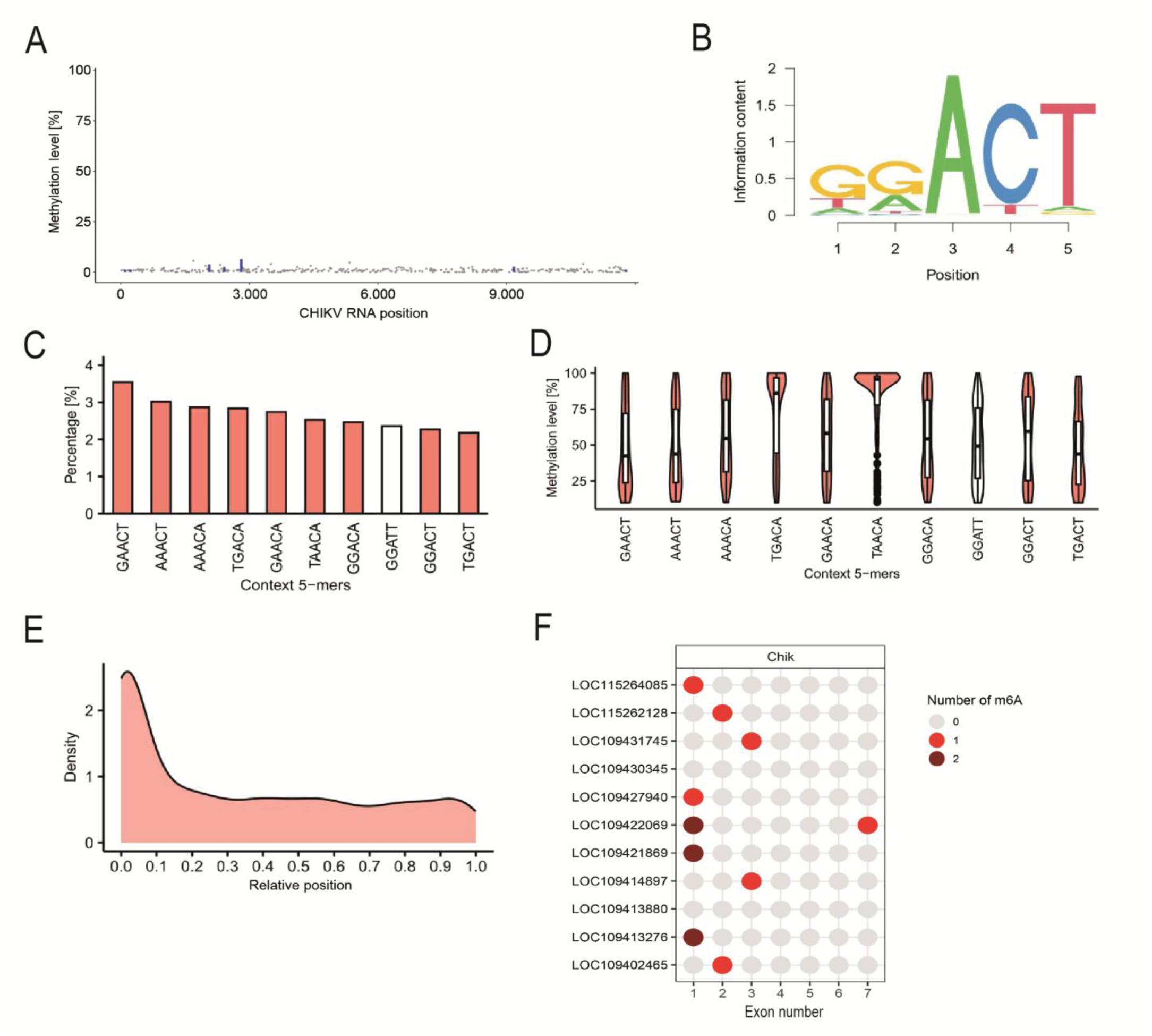
The m^6^A epitranscriptomic landscape of CHIKV-infected C6/36 cells. (A) GLORI-seq analysis of CHIKV viral RNAs. Grey dots indicate non-conversion levels of adenines in DRAC sites. Blue bars indicate potential methylation sites with a ratio >0.01 and Padj <0.05. Note that only a small fraction of sites exceeds these cutoffs and that their ratios are consistently low. (B) Sequence motif analyses using detected m^6^A sites in virus-infected C636 cells. m^6^A sites were primarily detected in the DRAC(H) consensus sequence motif. (C) Frequencies of the 10 most-detected motifs. (D) Quantification of the methylation level in the 10 most-detected motifs in virus-infected C636 cells. (E) Metagene plot depicting the positional distribution of detected m⁶A sites along normalized transcripts. m⁶A modifications are predominantly enriched within the CDS, with a higher density toward the 5′ region of the coding sequence. (F) Dotplot displaying distribution and number of detected m^6^A methylation sites in serine protease transcripts that were upregulated following pharmacological inhibition of METTL3 with STM2457, as identified by RNA-Seq. The results reveal that m^6^A modifications are broadly distributed throughout the CDS, without strong confinement to transcript termini, indicating preferential deposition within protein-coding regions under the analyzed conditions.

**Table 1.**
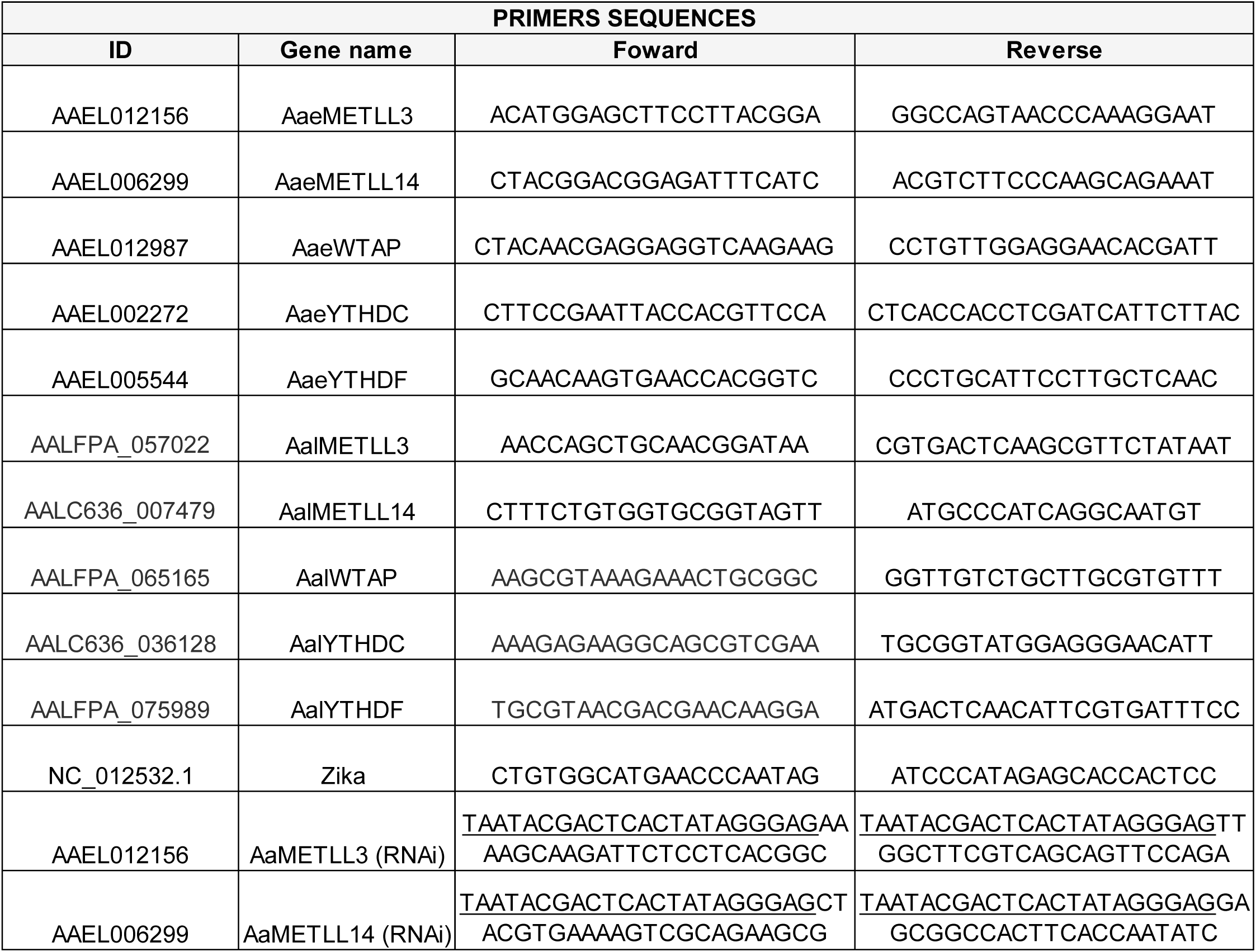
Gene names, GenBank accession numbers, and primer sequences used in this study. The table lists all target genes analyzed, including their corresponding GenBank accession IDs to ensure precise gene identification. Forward and reverse primer sequences (5′–3′) are provided for each gene. Primers were designed using Primer-BLAST (www.ncbi.nlm.nih.gov/tools/primer-blast and were validated prior to experimental use.

**Table 2.**
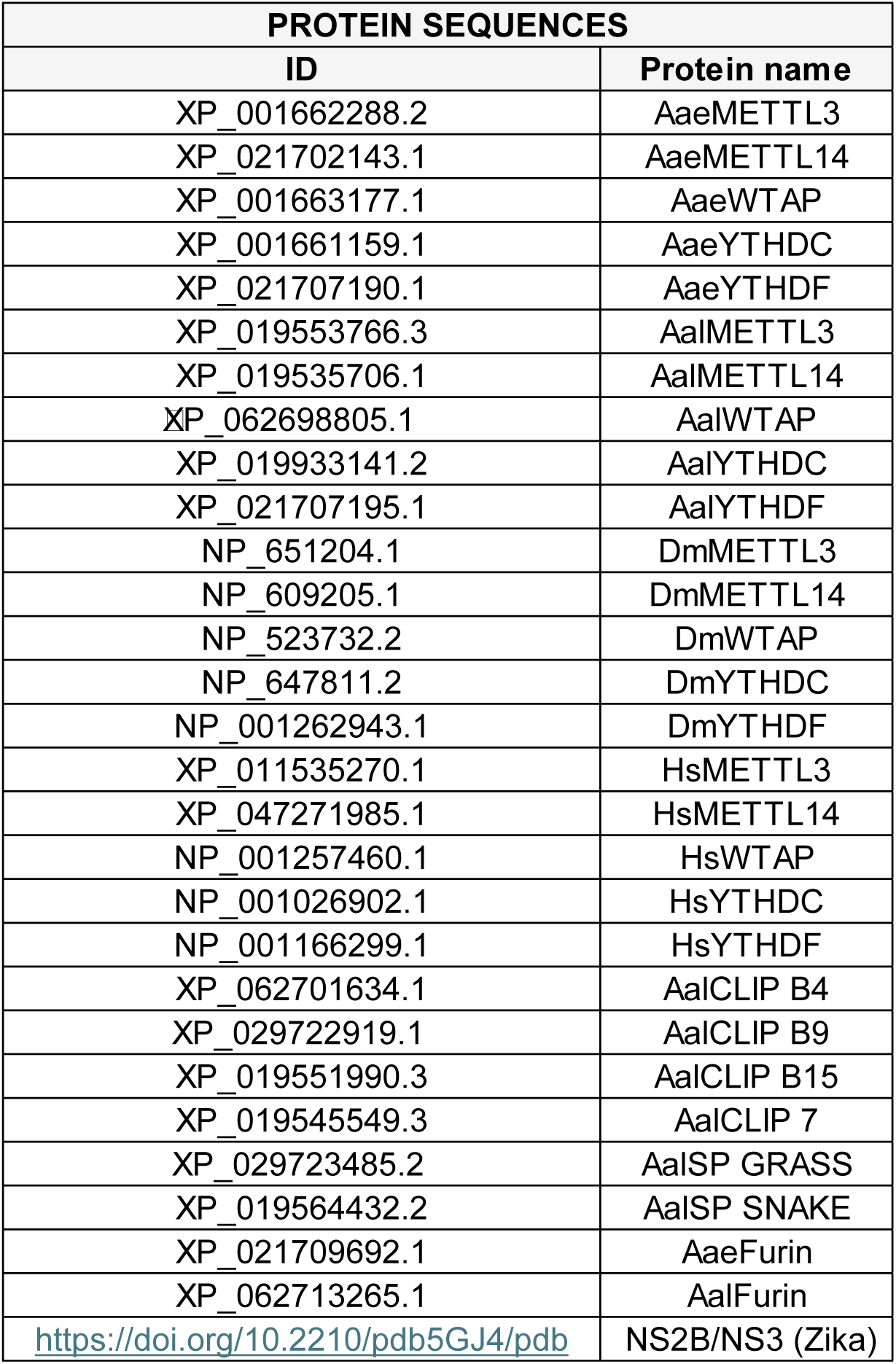
Protein names, GenBank accession numbers, and sequence information used for amino acid alignment and molecular docking analyses. The table lists all proteins included in the comparative analyses, together with their corresponding GenBank accession numbers to ensure unambiguous sequence identification. The indicated sequences were used to perform multiple sequence alignments for the assessment of conserved domains and catalytic residues. The protein sequences (or their derived structural models) were subsequently employed in molecular docking analyses to evaluate predicted ligand–protein interactions.

